# iPSC-derived skeletal muscle spheroids for Duchenne Muscular Dystrophy modeling

**DOI:** 10.1101/2025.06.05.657880

**Authors:** Joyce Esposito, Felipe de Souza Leite, Igor Neves Barbosa, Thaís Maria da Mata Martins, Giovanna Gonçalves de Oliveira Olberg, Ziad Al Tanoury, Kayque Alves Telles-Silva, Mayana Cristina da Silva Pardo, Tatiana Jazedje, Raul Hernandes Bortolin, Mario Hiroyuki Hirata, Olivier Pourquié, Mayana Zatz

**Affiliations:** Department of Genetics and Evolutionary Biology, Human Genome and Stem Cell Research Center, Biosciences Institute, University of São Paulo, São Paulo 05508-900, Brazil; Department of Clinical and Toxicological Analyses, School of Pharmaceutical Sciences, University of São Paulo, São Paulo 05508-000, Brazil; Department of Pathology, Brigham and Women’s Hospital, 60 Fenwood Road, Boston, MA 02115, USA; Division of Genetics and Genomics, Boston Children’s Hospital, Harvard Medical School, Boston, MA 02115, USA

**Keywords:** Muscle spheroids, induced pluripotent stem cells, Duchenne Muscular Dystrophy, Disease Modeling, Satellite Cells

## Abstract

**Background:** The progressive skeletal muscle degeneration observed in Duchenne Muscular Dystrophy (DMD) patients requires multiple cycles of satellite cells (SCs) activation to promote tissue regeneration. Dystrophic SCs present intrinsic defects, and the disrupting fibrotic niche hinders appropriate muscle recovery. Traditional 2D culture systems face challenges in modeling the DMD muscle niche and SCs behavior. Our aim was to validate a 3D culture of skeletal muscle spheroids (iSMS) for DMD modeling, as compared to the traditional 2D culture, while investigating the pathophysiological mechanisms of dystrophin deficiency *in vitro*.

**Methods:** To compare iSMS with traditional 2D myogenic differentiation, we differentiated PAX7 reporter wild-type (WT), dystrophic (DMD) isogenic induced pluripotent stem cells (iPSCs), and patients iPSCs, characterized myogenic markers levels and assessed differences in proliferation and differentiation using RT-qPCR, immunofluorescence, and flow cytometry.

**Results:** Our data showed that although both 2D and iSMS culture systems generated myogenic progenitors positive for *MYOD*, *MYOG*, *MYF5*, and *MYH3*, iSMS improved *PAX7* expression *in vitro*. Moreover, we identified a differential regulation of canonical Notch signaling genes between iSMS and 2D, which may influence the comparison between WT and DMD. We also characterized the differentiation of myogenic progenitors derived from 2D and iSMS towards elongated myofibers, providing a valuable comparison with muscle fibers differentiated from human primary myoblasts. Additionally, DMD iSMS derived progenitors proliferated at reduced levels compared with WT iSMS, a characteristic not observed in progenitors derived from 2D cultures. Finally, we performed iSMS and 2D myogenic differentiation of iPSC lines from three patients with DMD, thus validating the iSMS protocol for DMD modeling.

**Conclusion:** Our results highlight the important advantages of using the iSMS differentiation platform over 2D for disease modeling. Exploring these 3D systems may help to gain a deeper understanding of SCs behavior to advance in novel treatments for DMD, which might be applicable to other forms of muscular disorders.

## INTRODUCTION

Duchenne Muscular Dystrophy (DMD) is an X-linked genetic disease caused primarily by out-of-frame mutations in the *DMD* gene causing dystrophin deficiency (Hoffman et al., 1987; Monaco et al., 1988). In skeletal muscles, the lack of dystrophin leads to progressive degeneration and necrosis, extracellular matrix deposition, delayed regeneration, and consequent loss of muscular function over time (Emery, 2002). Previously considered a myofiber-only disorder, DMD has also been characterized by satellite cells (SCs) dysfunction and consequent impaired regenerative capacity (Kodippili & Rudnicki, 2023).

SCs are formed in the late fetal stage, and in adult muscles, they are characterized by Pax7 expression (Seale et al., 2000; Picard & Marcelle, 2013). In homeostasis, SCs are maintained in a quiescent state closely associated with myofibers, between the sarcolemma and basal lamina (Zhou et al., 2022). After muscle injury, SCs are activated and enter the cell cycle, which is accompanied by the temporal and spatial expression of the myogenic regulatory factors Myf5, MyoD, MyoG, and MRF4, a family of basic helix-loop-helix transcription factors responsible for regulation of prenatal and postnatal myogenesis (Zammit, 2017). The balance between SCs activation and quiescence and between asymmetric and symmetric cell divisions determines the maintenance of the stem cell pool throughout life (Ancel et al., 2021). However, this balance is severely compromised in dystrophin-deficient muscles due to constant regeneration and degeneration cycles accompanied by SCs intrinsic defects and the disrupting niche (Kodippili & Rudnicki, 2023).

Studying skeletal muscles and the dynamics of SCs formation and maintenance in DMD has been challenging because of difficulties in extrapolating animal model results to human disease contexts (Wells, 2018). The *in vitro* culture of human cells can circumvent these challenges, but conventional bidimensional (2D) cultures fail to reproduce critical aspects of muscle physiology, stem cell fate, and niche, hindering the maintenance of SCs for long periods (Montarras et al., 2005; Pang et al., 2023). Furthermore, ethical and technical issues limit isolation and expansion of SCs from the skeletal muscle of DMD patients (Zschüntzsch et al., 2022).

Numerous studies have focused on the development of advanced 3D skeletal muscle models that better replicate skeletal muscle interactions, mechanical loading, stiffness, and extracellular matrix composition (Jalal et al., 2021). These models may also improve our knowledge of the pathological mechanisms of neuromuscular diseases, contributing to drug development and precision medicine (Khodabukus et al., 2018). However, only a limited number of studies have reported the generation and maintenance of human SCs in these culture systems, which is a current challenge in the field, especially in the context of DMD.

The advent of induced pluripotent stem cells (iPSCs) has enabled possible the investigation of patient-specific cells and the molecular mechanisms governing disease progression (Takahashi et al., 2007; Danisovic et al., 2018). Transgene-free protocols for the myogenic differentiation of human iPSCs can recapitulate key aspects of early skeletal muscle development, ultimately giving rise to PAX7^+^ cells with SCs properties (Chal et al., 2015; Tanoury et al., 2020). Despite being a valuable model for studying patient-specific SCs and myogenic progenitors, there are challenges in establishing appropriate phenotypic *in vitro* parameters to compare healthy and dystrophic cells (Shoji et al., 2015; Choi et al., 2016). However, the poor complexity of traditional 2D cultures can be improved using more physiological 3D models (Piga et al., 2019). For instance, artificial 3D hydrogel-based muscles from iPSCs of patients with skeletal muscle laminopathies highlighted nuclear abnormalities and deformities that were not apparent in conventional 2D cultures (Maffioletti et al., 2018).

In this study, we have performed differentiation of iPSCs into skeletal muscle spheroids (iSMS). Our aim was to produce myogenic progenitors and SC-like cells from iPSCs, comparing iSMS with conventional 2D culture using a DMD lineage (DMD) and its parental isogenic WT lineage. We showed a similar differentiation capacity in 2D and iSMS from both WT and DMD lineages, but a marked increase in *PAX7* mRNA expression in iSMS. Furthermore, we characterized Notch pathway regulation and proliferative behavior in 2D and iSMS from WT and DMD isogenic lineages. We also provided a valuable comparison between skeletal muscle fibers derived from iPSCs and primary human adult myoblasts. Finally, we characterized iSMS from patient-derived iPSCs with different pathogenic variants of *DMD*.

## RESULTS

### Differentiation into myogenic progenitors in 2D versus iSMS culture systems

First, we tested whether myogenic differentiation could be performed to generate 3D iSMS using the protocol described by Chal et al. (2016) for adherent cultures, and potential outcome differences between WT and DMD isogenic lines. Therefore, 2D and iSMS differentiation of iPSCs into myogenic progenitors was performed side by side **(Figure 1a)**. No visible morphological changes were observed in the 2D cultures comparing DMD and WT **(Figure 1b)**, and DMD and WT iSMS displayed similar dimensions **(Figure 1c)**.

**Figure 1.**
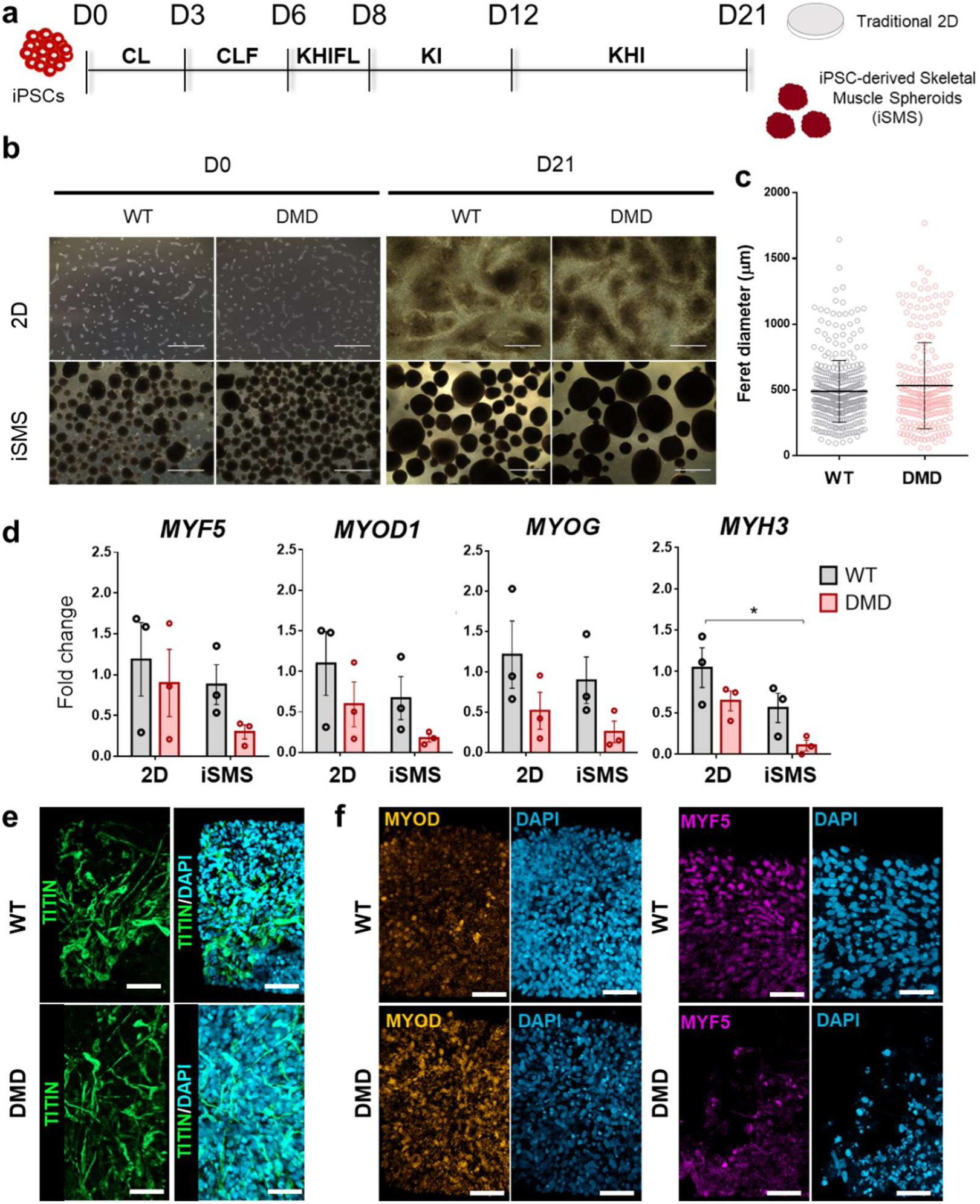
Skeletal muscle differentiation of WT and DMD iPSCs in iSMS versus traditional 2D cultures. **a.** Schedule of primary differentiation of iPSCs to the myogenic lineage and formation of skeletal muscle spheroids (iSMS) or traditional 2D differentiation. The medium composition in each phase is distinguished by the following letters for the main reagents/factors used: C stands for CHIR-99021; L, LDN-193189; F, FGF2; K, KSR; H, HGF; and I, IGF. **b.** Representative images of cells from the WT and DMD lineages on day 0 (D0) and on the final day of primary differentiation (day 21, D21) in 2D and iSMS. Scale: 1000 µm. **c.** Evaluation of spheroids size from WT and DMD (n=226 iSMS for DMD and n=361 for WT). **d.** Gene expression analysis of *MYOD1, MYF5, MYOG* and *MYH3* in 2D and iSMS cultures of the WT and DMD lineages (n=3). WT 2D was used as a reference (fold-change = 1). We performed twoway ANOVA and Tukey’s multiple comparisons test to check for statistical significance (* represents p<0.05). **e.** Immunostaining of iSMS from WT and DMD on D21 showing cells positive for TITIN (green). Cell nuclei are shown in blue. **f.** Immunostaining of iSMS from WT and DMD on D21 showing positive staining for MYOD (orange) and MYF5 (purple). Cell nuclei are shown in blue. Scale E and F: 50 µm.

Gene expression analysis confirmed the expression of the myogenic markers *MYOD, MYOG, MYF5,* and *MYH3* in both 2D and iSMS, thus validating the use of the original protocol to generate iSMS **(Figure 1d)**. Overall, these markers were not statistically different when comparing 2D versus iSMS or WT versus DMD. Moreover, both WT and DMD iSMS generated TITIN, MYOD, and MYF5 positive cells **(Figure 1e and 1f)**.

### Derivation of PAX7^+^ SC-like cells in differentiated WT and DMD in 2D and iSMS

We hypothesized whether iSMS provide a more suitable environment for the generation and maintenance of PAX7^+^ cells during differentiation. Also, we investigated whether DMD would differ from WT in terms of SCs formation and maintenance. On differentiation day 21, a Venus YFP signal was detected on iSMS **(Figure 2a)** which was confirmed by immunocytochemistry for PAX7 **(Figure 2b)**. While flow cytometry revealed a similar percentage of PAX7^+^ cells for WT and DMD in both 2D and iSMS **(Figure 2c and 2d)**, the *PAX7* mRNA level was almost 10 times higher in WT iSMS versus WT 2D, and in DMD iSMS compared to DMD 2D **(Figure 2e)**. Interestingly, while *PAX7* mRNA was significantly reduced in DMD iSMS versus WT iSMS, there were not significant differences between WT and DMD in 2D cultures. These data indicated that iSMS may be more suitable for the derivation and maintenance of PAX7^+^ cells from iPSCs at this point of myogenic differentiation.

**Figure 2.**
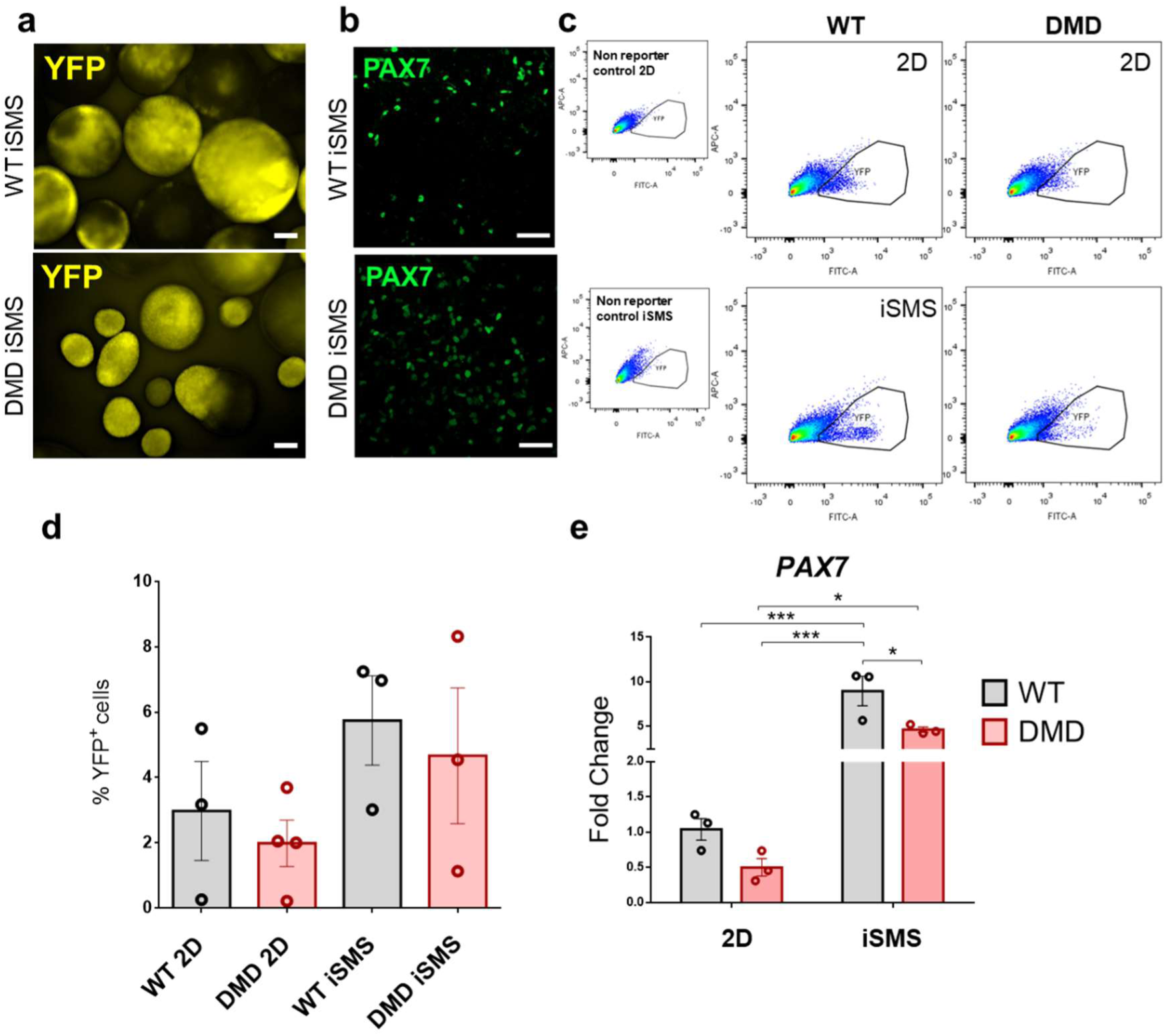
PAX7 expression in iSMS and 2D cultures after *in vitro* myogenic differentiation. **a.** Identification of YFP positive cells in iSMS after differentiation of WT and DMD iPSC YFP reporter lines. Scale: 250 µm. **b.** Immunofluorescence of WT and DMD iSMS for identification of PAX7 positive cells (green). Scale: 50 µm. **c.** Flow cytometry results show the presence of YFP^+^ cells in 2D and iSMS cultures. **d.** The proportion of YFP^+^ cells in WT and DMD 2D and iSMS obtained from flow cytometry data (n=at least 3 independent experiments). **e.** Results of *PAX7* mRNA expression on day 21 comparing 2D and iSMS of WT and DMD groups (n=3). WT 2D was used as a reference (foldchange=1). Statistical significance was evaluated by TwoWay ANOVA and Tukey’s multiple comparisons (* represents p<0.05, and *** p<0.001).

### Notch pathway regulation in iSMS and 2D cultures

The Notch pathway is a major regulator of quiescence and SC fate acting as a cell-cell communication system that might benefit from the iSMS 3D environment. Thus, we hypothesized that the Notch pathway would be differentially regulated in iSMS as compared to traditional 2D cultures, especially when comparing WT and DMD. Thus, we performed an RT^2^ profiler array for Notch pathway genes and investigated their expression profiles in 2D and iSMS from WT and DMD. Of the 84 Notch pathway target genes **(Supplementary Table 3)**, 13 were excluded because they were not expressed in detectable amounts in the samples.

We also compared WT with DMD in both 2D and iSMS, aiming to explore possible differences between the two culture types. When we compared DMD with WT in 2D cultures, we found 16 downregulated genes, and 19 upregulated genes. On the other hand, in iSMS we found 41 downregulated genes and 5 upregulated genes, suggesting a greater reduction in Notch pathway regulation in DMD iSMS than in 2D **(Figure 3a)**. We found 24 genes that were commonly differentially expressed in both 2D and iSMS when comparing WT and DMD.

**Figure 3.**
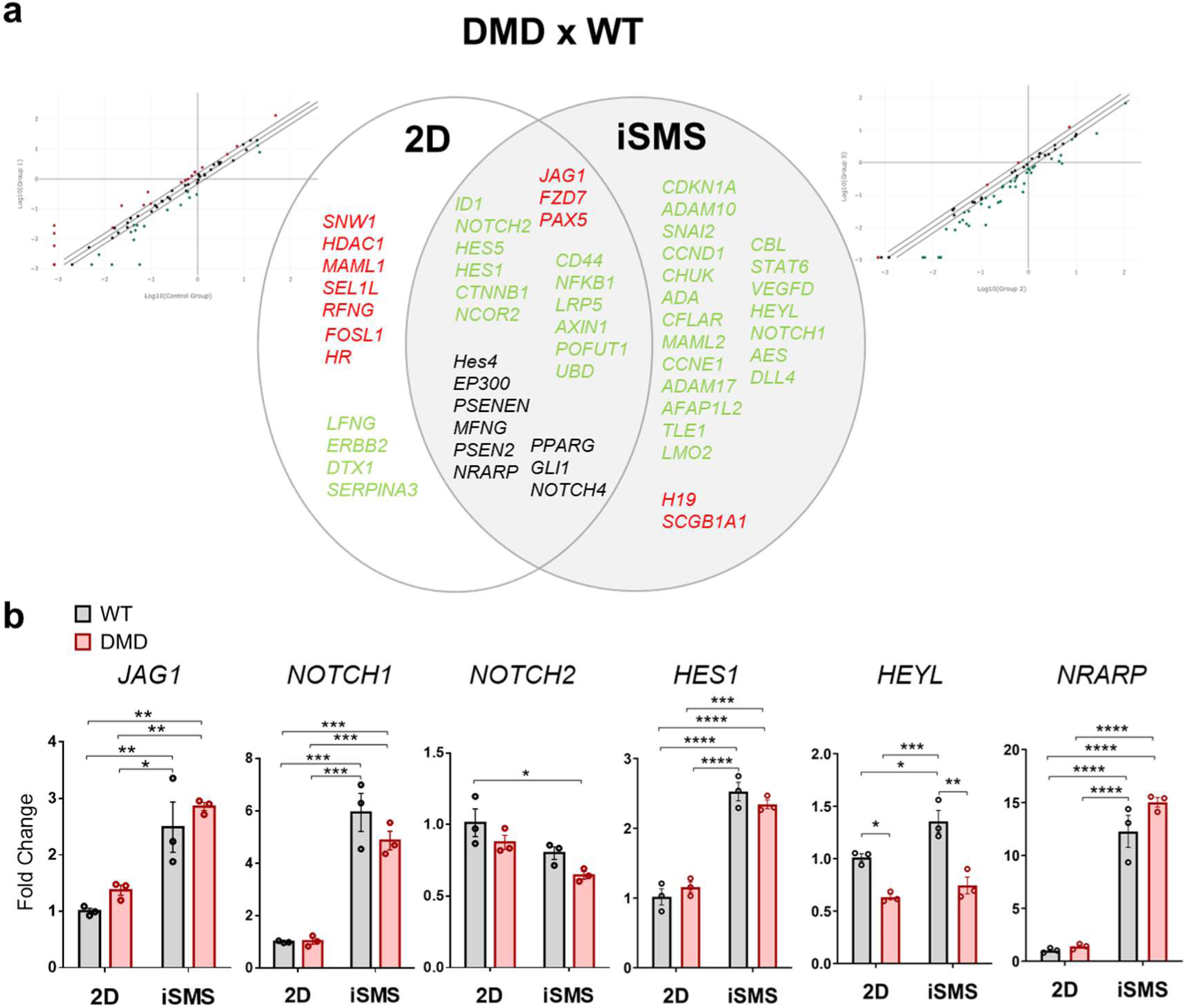
Notch pathway modulation in 2D versus iSMS from WT and DMD lineages. **a.** Results of Notch pathway RT^2^ PCR array showing genes that are modulated comparing WT and DMD in 2D and iSMS. Scatter plot showing differentially expressed genes obtained from 2D (left) and iSMS (right). Center: Venn diagram showing DEGs. In green and red are represented possibly downregulated or upregulated genes, respectively, in DMD versus WT within the correspondent culture type. In black are the genes that were upregulated in one group and downregulated in another group, comparing DMD to WT, n=1. **b.** Gene expression analysis of *JAG1*, *NOTCH1*, *NOTCH2*, *HES1*, *HEYL*, and *NRARP* mRNA. WT 2D was used as a reference (foldchange=1). Statistical significance was evaluated by TwoWay ANOVA and Tukey’s multiple comparisons (* represents p<0.05, ** p<0.01, *** p<0.001, and **** p<0.0001).

Focusing on key canonical Notch pathway components, we analyzed their gene expression patterns using RT-qPCR. Compared to WT 2D, WT iSMS exhibited upregulation of *JAG1*, *NOTCH1*, *HES1*, *HEYL* and *NRARP*, but not *NOTCH2* **(Figure 3b)**. Similarly, DMD iSMS showed increased expression of *JAG1*, *NOTCH1*, *HES1* and *NRARP*, but not *NOTCH2* and *HEYL*, as compared to DMD 2D. Notably, when comparing DMD and WT, *HEYL* was the only gene significantly downregulated in the DMD group under both 2D and iSMS conditions. Our findings highlight that the type of cell culture may directly influence Notch signaling in myogenic progenitors from iPSCs.

### Secondary muscle differentiation in 2D and iSMS cultures

Next, we investigated whether myogenic progenitors in iSMS could be further differentiated (secondary differentiation) into skeletal muscle fibers. For 2D cultures, this step is preceded by cell dissociation, which is a harsh procedure, and upon culture with myoblast growth medium, selectively favors myogenic progenitors proliferation, followed by differentiation into skeletal muscle fibers **(Figure 4a)**. As iSMS was not subjected to dissociation and replating, it is important to mention that other cell types would still be present in iSMS. Despite this, we hypothesized that iSMS would maintain a higher *PAX7* expression than 2D cultures. The 2D and iSMS cultures were evaluated after 8 days of secondary differentiation. Importantly, the terminal differentiation medium contains SB-431542, a potent TGF-B receptor type I kinase inhibitor that stimulate myotube fusion (Hicks et al., 2018).

**Figure 4.**
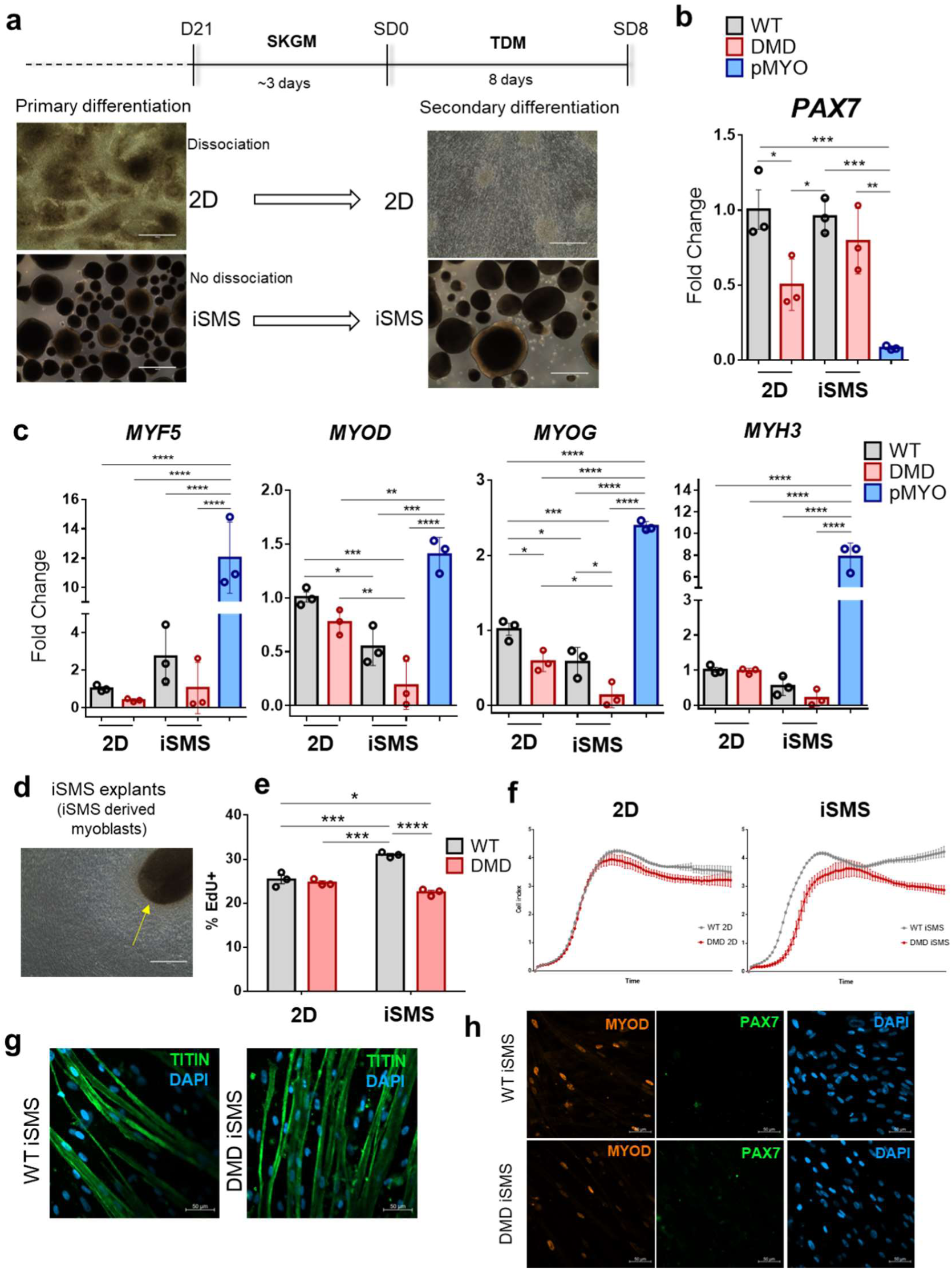
Secondary myogenic differentiation of iPSCs in 2D and iSMS and comparison with pMYO. Schedule of secondary myogenic differentiation of 2D and iSMS cultures. After primary differentiation, cells were transferred to SKGM-2 medium for 3 days and then cultivated in terminal differentiation medium (TDM) for 8 days for derivation of skeletal muscle fibers. **b** and **c.** Gene expression of *PAX7*, *MYF5*, *MYOD, MYOG* and *MYH3* comparing 2D and iSMS culture systems of WT and DMD lineages and primary control CD56^+^/CD82^+^ myoblasts (pMYO) differentiated into skeletal muscle fibers. WT 2D was used as a reference, and RPLP0 was utilized for normalization using ΔΔCT method. One-way ANOVA and Tukey’s multiple comparisons test were performed to check statistical significance. * indicates p<0.05, ** p<0.01, *** p<0.001, and **** p<0.0001. **d.** Representative image of migrating cells from iSMS explants when maintained in Matrigel coated plates. **e.** EdU Cell proliferation assay results showing the percentage of proliferative cells of iPSC-derived myogenic progenitors of WT and DMD from 2D and iSMS. **f.** Proliferation assay of iPSC-derived myogenic progenitors of WT and DMD in 2D and iSMS using xCELL ligence real-time cell analyzer. **g.** Representative immunostaining skeletal muscle fibers from iSMS-derived progenitors showing TITIN positive elongated fibers. **h.** Immunostaining for MYOD (orange) and PAX7 (green) of skeletal muscle fibers from iSMS-derived progenitors. In g and h nuclei are represented in blue (DAPI). Scale: 50 µm.

*PAX7* mRNA was detected in all groups after secondary differentiation, and expression levels were similar between WT 2D and WT iSMS, and between DMD 2D and DMD iSMS **(Figure 4b)**. Contrary to what we observed in primary differentiation, *PAX7* was reduced in DMD 2D versus WT 2D, but this difference was not found in WT iSMS versus DMD iSMS. To better investigate this, we analyzed *PAX7* expression by comparing the data from primary differentiation day 21 (D21) with the secondary differentiation day 8 (SD8). This analysis revealed marked *PAX7* downregulation in iSMS after secondary differentiation compared to D21, while no changes were observed in 2D cultures **(Supplementary Figure 1)**. These data revealed that the peak of *PAX7* expression in iSMS was obtained during primary differentiation, suggesting that this is an optimum time point for studying PAX7^+^ SC-like cells.

Additionally, we compared differentiated myogenic progenitors from iPSCs with differentiated CD56^+^/CD82^+^ human primary myoblasts (pMYO), a highly myogenic population, purified as previously described (Spinazzola & Gussoni, 2017) **(Supplementary Methods and Supplementary Figure 2)**. *PAX7* expression was markedly reduced in pMYO compared to iPSC-derived skeletal muscles **(Figure 4b)**. The expression levels of *MYF5* and *MYH3* were not different among the groups, except when compared with pMYO **(Figure 4c)**. *MYOD* was downregulated in iSMS, as compared to 2D cultures from WT and DMD, and upregulated in pMYO. Consistently, the expression of *MYOG*, a terminal differentiation marker, was reduced in DMD compared to WT in both 2D and iSMS cultures. The expression levels of *MYF5*, *MYOD*, *MYOG* and *MYH3* were higher in pMYO, revealing differences in the maturity of iPSC-derived myotubes versus adult pMYO-derived myotubes.

### Expansion of myogenic progenitors from iSMS

We next asked whether SC generated in iSMS could be dissociated and expanded in a 2D environment and still derive skeletal muscle fibers. After SD8, the iSMS were transferred to Matrigel-coated plates. The iSMS grew in SKGM-2 medium, and the cells adhered to and migrated out of them **(Figure 4d)**.

The migrating myogenic progenitors from iSMS and 2D-derived myogenic progenitors were subcultured and frozen for long-term storage. After thawing and expanding in SKGM-2 medium, we compared cell proliferation and maintenance of the differentiation capacity of both 2D and iSMS culture-derived myoblasts from WT and DMD. We found a higher percentage of EdU-positive cells in the myogenic progenitors derived from WT iSMS than all other groups, especially comparing with myogenic progenitors from DMD iSMS **(Figure 4e)**. In contrast, the percentage of proliferative myogenic cells did not differ between the WT 2D and DMD 2D. Proliferation curves using xCELLigence real-time analysis for 18 days confirmed the higher differences between WT and DMD in cell proliferation of myogenic progenitors derived from iSMS **(Figure 4f).** These data suggest that iSMS is a suitable model for evaluating the consequences of *DMD* pathogenic mutations on the proliferative behavior of iPSC-derived myogenic progenitors.

We confirmed the differentiation capacity of myogenic progenitors derived from the iSMS protocol after secondary differentiation and freezing/thawing cycle, similar to what was reported for 2D cultures (Chal et al., 2015). Immunocytochemistry with antibodies against TITIN **(Figure 4g)** confirmed the differentiation of WT and DMD iSMS myogenic progenitors into iPSC-derived myofibers. Additionally, we observed the presence of MYOD-positive cells, whereas PAX7-positive cells were nearly absent at this point **(Figure 4h)**.

### 3D iSMS versus 2D differentiation from DMD patient derived iPSCs

We also aimed to assess how iSMS differentiation occurs in iPSCs from unrelated DMD patients. The iPSCs were produced and characterized accordingly and expressed pluripotent stem cell markers, as confirmed by RT-qPCR of *SOX2*, *NANOG*, and *OCT4* **(Supplementary Figure 3a)**, and immunofluorescence for OCT4 and SSEA4 **(Supplementary Figure 3d)**. We also confirmed that iPSCs were free of chromosomal imbalances **(Supplementary Figure 3b and 3c)**.

We performed 2D and iSMS differentiation of three iPSC lineages from DMD patients (DMD-A, DMD-B and DMD-C; **Supplementary Table 2**), and one control (CTRL-1). To characterize the differences between the corresponding 2D differentiation, we identified the PAX7 protein and evaluated the expression of myogenic markers. As observed for the isogenic cell lines, PAX7 positive cells were present in both 2D and iSMS at similar percentages **(Figure 5a and Figure 5b).** Interestingly, when combining the CTRL-1, and DMD patient groups, we observed a higher median fluorescence intensity (MFI ratio) in iSMS, than in 2D **(Figure 5c)**.

**Figure 5.**
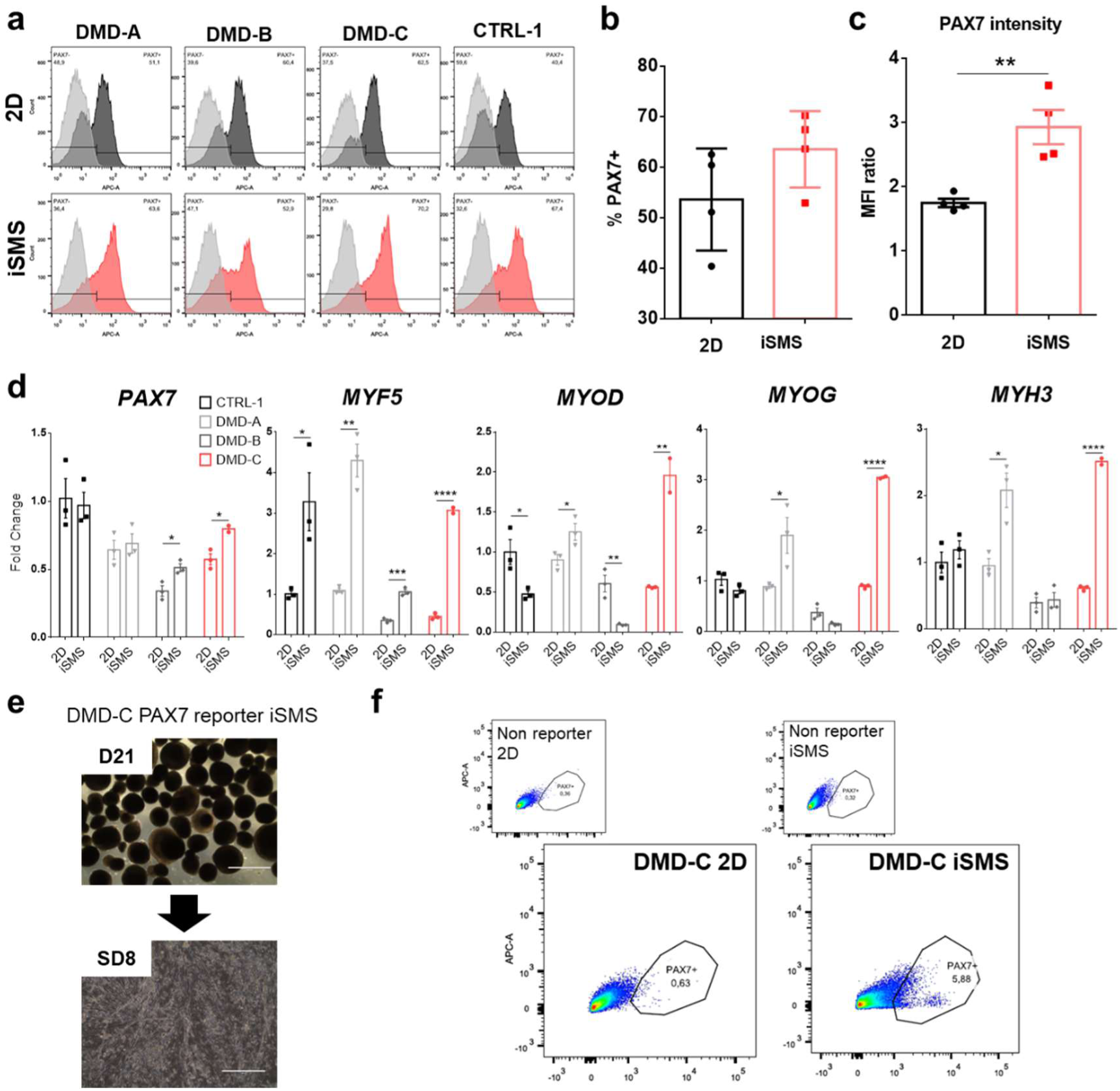
Myogenic differentiation of iPSCs from DMD patients in 2D versus iSMS. **a.**Flow cytometry results of *PAX7* expression indicating SC-like cells formation in both cultures. **b.** Quantification of the proportion of PAX7^+^ cells in 2D and iSMS from flow cytometry data presented in a. These data include CTRL-1, DMD-A, DMD-B, and DMD-C levels in the same group. **c.** Quantification of PAX7 intensity (Mean Fluorescence Intensity ratio) from the flow cytometry data of 2D and iSMS cultures. These data include CTRL-1, DMD-A, DMD-B and DMD-C in the same group. Unpaired t tests were performed, and ** indicates p<0.01. **d.** Gene expression analysis of *PAX7*, *MYF5*, *MYOD*, *MYOG*, and *MYH3*. Data are represented as mean ± SEM. An unpaired t test was performed to compare each lineage in 2D with the corresponding iSMS. * represents p<0.05, ** p<0.01, *** p<0.001, and **** p<0.0001. CTRL-1 2D was used as a reference (foldchange = 1), and *RPLP0* was applied for normalization using the ΔΔCT method. **e.** Representative images of DMD-C PAX7 reporter cells after primary (D21) and secondary (SD8) myogenic differentiation. **e.** Flow cytometry of Venus-YFP from 2D and iSMS cultures of DMD-C.

Regarding gene expression, we compared 2D cultures with iSMS from the same patient to evaluate the corresponding effect of the 3D culture system **(Figure 5d)**. In DMD-C, iSMS were shown to upregulate all evaluated myogenic markers *PAX7*, *MYF5, MYOD, MYOG* and *MYH3*, comparing to 2D. The iSMS from DMD-A also showed an upregulation of all myogenic markers, except *PAX7*, compared to its corresponding 2D. CTRL-1 iSMS demonstrated an upregulation of *MYF5*, downregulation of *MYOD*, and a similar level of other myogenic markers comparing to 2D. Surprisingly, iSMS from DMD-B showed an upregulation of *PAX7* and *MYF5* mRNA, but exhibited downregulation of *MYOD*, compared to 2D. These data reinforces the idea that patient-specific iPSCs may exhibit heterogeneous behavior during *in vitro* myogenic differentiation, as previously described (Choi et al., 2016). Despite their variable efficiencies, it is possible to perform myogenic induction from DMD patient-derived iPSCs, and iSMS may be a good platform to study DMD.

To further validate the feasibility of this 3D model for the derivation of PAX7^+^ SC-like cells from iPSCs, we generated a PAX7 reporter iPSC from patient DMD-C, since this cell line displayed an increase in the expression of all evaluated myogenic markers comparing 2D with iSMS. We showed that progenitors derived from the iSMS reporter DMD-C line were able to differentiate after subculturing **(Figure 5e)** and, in this case, generated more PAX7^+^ cells than 2D culture **(Figure 5f)**. This emphasizes the relevance of 3D culture in generating SC-like cells from iPSCs from dystrophic patients. Overall, this could be particularly important in the study of DMD, since dystrophin deficiency directly affects SCs fate (Dumont et al., 2015).

## DISCUSSION

Modeling genetic diseases with iPSCs has been an exciting approach to uncover the molecular and cellular mechanisms controlling patients’ phenotype (Takahashi et al., 2007; Liu et al., 2018). The iPSCs-derived progenitors can successfully recapitulate key aspects of diseases as previously shown for amyotrophic lateral sclerosis (Oliveira et al., 2020), Parkinson’s disease (Nguyen et al., 2011), Down syndrome (Kawatani et al., 2021), Alzheimer’s disease (Yagi et al., 2011), DMD (Danisovic et al., 2018), among many others. However, the use of iPSC technology to model muscle diseases is challenging due to poor reproducibility and efficiency, distinct genetic backgrounds, lack of appropriate natural cell-cell and cell-matrix interactions, and of mechanical and physical properties (Dessauge et al., 2021; Chien et al., 2022). To overcome some of these limitations, an increasing number of studies have aimed to develop *in vitro* 3D skeletal muscles models using pluripotent stem cells (Hosoyama et al., 2014; Sakai-Takemura et al., 2018; Maffioletti et al., 2018; Rao et al., 2018; Mavrommatis et al., 2023).

DMD is the most severe form of muscular dystrophy and is characterized by progressive skeletal muscle degeneration and SC dysfunction over time (Duan et al., 2021). Understanding SC behavior provide new perspectives for the treatment of this devastating disease (Filippelli & Chang, 2021). Here, we generated iPSCs-derived muscle spheroids (iSMS) adapted from previously described transgene-free (Chal et al., 2015, 2016) and free-floating culture (Hosoyama et al., 2014) protocols. Our main question was: Is iSMS a more suitable culture system than 2D for DMD *in vitro* modeling?

Three weeks were sufficient to derive PAX7^+^ SC-like cells, as evidenced by the presence of YFP^+^ cells and the identification of PAX7 protein and mRNA. This time point was chosen based on previous results indicating that *PAX7* peaked at approximately at day 21 of primary differentiation (Tanoury et al., 2020). Additionally, floating cultures for long periods (over six weeks) may result in reduced myogenic activity (Sakai-Takemura et al., 2018). Our data revealed that *DMD* out-of-frame mutation in the isogenic iPSC DMD did not seem to significantly compromise the expression of the myogenic markers *MYOD*, *MYF5*, *MYOG* and *MYH3* during primary skeletal muscle differentiation in 2D and iSMS. On the other hand, it had a major impact on *PAX7* mRNA expression, which was evident when comparing iSMS, but not in 2D culture. It suggests that dystrophic iSMS (DMD) may present intrinsic defects in upregulating or maintaining *PAX7* expression during primary myogenic differentiation, compared to its isogenic WT.

Although we observed this difference in *PAX7* expression, the proportion of SCs in iSMS was similar between WT and DMD. Correspondingly, a previous study employing 2D culture of this DMD iPSC lineage showed no significant differences between WT and DMD in the number of PAX7 positive cells (Tanoury et al., 2021). Considering our gene expression data, one hypothesis is that, individually, WT SC-like cells express higher levels of *PAX7* (possibly forming more PAX7^High^ cells) than DMD SC-like cells. Overall, despite being distinct models, this is in agreement with previous studies showing that dystrophin deficient SCs lose quiescence precociously, whereas the number of SCs is maintained or elevated in dystrophic muscles (Kottlors & Kirschner, 2010; Ribeiro et al., 2019).

When secondary differentiation was performed, a more distinct gene expression pattern was observed: *PAX7* was downregulated in DMD 2D as compared to WT 2D, but not in WT iSMS versus DMD iSMS. We hypothesized that this is because the terminal differentiation medium (TDM) strikingly reduced *PAX7* mRNA levels in iSMS, compared with primary differentiation. In fact, *PAX7* downregulation is expected and has been reported after long-term differentiation of iPSCs (Hosoyama et al., 2014; Jiwlawat et al., 2017), since TDM promotes the activation of SC-like cells to fuse with existing fibers. Accordingly, a previous study showed that the percentage of quiescent PAX7^+^/Ki67^-^ cells in skeletal muscle organoids was higher on day 30 than in subsequent time points day 70 and 130 (Shin et al., 2022). Furthermore, it is known that SB-431542 treatment accelerates cell fusion (Hicks et al., 2018).

Our data suggest that D21 of primary differentiation is a more appropriate time point for the analysis or isolation of PAX7^+^ SC-like cells produced in the iSMS. Improving the efficiency of PAX7 positive cells production is particularly important for isolating this population for downstream applications. For example, FACS sorting of a small subset of weakly stained positive cells may be challenging and may result in low efficacy. On the other hand, sorting PAX7 reporter iPSC cells from iSMS could be significantly more straightforward. Selection of the PAX7^+^ population may be necessary because of the intrinsic heterogeneity and technical issues of myogenic differentiation protocols, which can derive multiple other cell types, such as mesenchymal cells and neuroprogenitors (Xi et al., 2020). Furthermore, SCs are a naturally heterogeneous population (Tierney & Sacco, 2016), and understanding the cellular and molecular mechanisms that determine the different SC subpopulations is of great relevance for the development of therapies to prevent progressive muscle degeneration in DMD.

Comparing 2D versus iSMS after secondary differentiation, in both WT and DMD groups, a reduction in *MYOD* and *MYOG* levels suggests a more quiescent inactive state in myogenic progenitors from iSMS, although further characterization is required. Accordingly, myospheres formed by muscle stem cells from mice can be sustained in a pre-myogenic state in culture; upon stimulation, they start to proliferate, upregulate MyoD expression and can be differentiated into multinucleated myotubes (Westerman et al., 2010).

Notch signaling is a highly conserved pathway essential for SCs self-renewal, differentiation, and quiescence, being essential for determining SCs homing in their niche (Bjornson et al., 2012; Bröhl et al., 2012; Mourikis et al., 2012). Through cell-cell interactions, the transmembrane Notch receptors Notch1, Notch2, Notch3 or Notch4 interact with Notch ligands Dll1, Dll3, Dll4, Jag1 or Jag2, thereby activating intracellular molecular events culminating with the expression of the transcription factors of the Hes and Hey family (Zhou B. et al., 2022). As Notch signaling depends on cell-cell interaction and directly dictate *PAX7* expression (Wen et al., 2012), we investigated how the type of culture would modulate this pathway. In an exploratory analysis, we found common DEGs between WT and DMD in iSMS and 2D. Most importantly, our data showed that a greater number of Notch related genes were downregulated in DMD iSMS as compared to WT iSMS than the same comparison in 2D. It is well established that diminished Notch signaling contributes to DMD pathogenic mechanisms (Hartog & Asakura, 2022). For example, Notch signaling blockade in mouse SCs led to muscular dystrophy phenotype and impaired muscle regeneration (Lin et al., 2013). A missense variant in *POGLUT1* gene causes autosomal recessive limb-girdle muscular dystrophy with decreased levels of Notch activation and Notch-dependent loss of SCs (Servián-Morilla et al., 2016). When looking for specific canonical Notch components, we noted an upregulation as a consequence of the iSMS culture, in both WT and DMD, suggesting that the Notch pathway may be strongly activated. Our results reveal that iSMS is a good platform for studying DMD and Notch pathway, paving the way for future studies modulating Notch for therapeutic purposes.

In this study, we also identified that myogenic progenitors from isogenic DMD iSMS displayed reduced cell proliferation, as evidenced by EdU and real-time xCELLigence experiments, whereas no differences were observed in 2D DMD. Several studies have reported proliferative defects in dystrophic primary myoblasts and replicative senescence due to telomere shortening (Blau et al., 1983; Webster & Blau, 1990; Sacco et al., 2010). Escaper GRMD dogs overexpressing *Jagged1* displayed greater proliferative capacity (Vieira et al., 2015). Conversely, other study showed that dystrophic myoblasts exhibited increased proliferation (Gosselin et al., 2022).

Here, we validated the generation of iSMS using iPSCs from DMD patients. Successful differentiation was achieved. Resembling what we observed for WT and DMD isogenic lines, we noted a similar proportion of PAX7^+^ cells, but higher *PAX7* levels in iSMS versus 2D, although with diverse expression pattern of myogenic markers. Concordantly, a study using a similar sphere-based myogenic differentiation protocol described different efficacies of four iPSC lines from different donors (Sakai-Takemura et al., 2018). Despite the heterogeneity observed, we showed the feasibility of producing iSMS from patients iPSCs independent of the *DMD* mutation. Strategies for selecting the PAX7^+^ subset, – for example using the DMD-C PAX7 reporter lineage - or single cell RNA sequencing may be required to increase the robustness of the comparisons.

Typically, most protocols generate cells by using 2D cultures before transitioning them into a 3D context (Jalal et al., 2021). In this study, we produced myogenic progenitors directly in a 3D environment and demonstrated that they maintained their proliferation and differentiation potential. Our primary differentiation did not involve Matrigel or any type of xenogenic extracellular matrix component, favoring their therapeutic application, since Matrigel exhibits batch-to-batch variability and may contain xenogeneic contaminants (Aisenbrey & Murphy, 2020). In addition, the originally described sphere-based protocol require spheres dissociation weekly (Hosoyama et al., 2014), and we speculate that this could disturb the quiescence/activation dynamics of SC-like cells during differentiation. Our adapted iSMS protocol has the potential to generate myogenic progenitors for use in cell therapy and in the development of 3D muscle constructs for disease modeling.

Overall, our data suggests that iSMS is a good platform for studying human DMD, opening new perspectives to understand SCs disease-related dysfunction and to develop novel therapeutic strategies for this devastating muscle disease.

## METHODS

### iPSC Lines and Patients

CRO wild type (WT) and CRO DMD (DMD) isogenic cell lines have been produced and characterized elsewhere (Tanoury et al., 2020, 2021). We introduced a knock-in in exon 1 of *PAX7* promoter in the WT and DMD lineages to create PAX7-Venus reporter lineages, as previously described (Tanoury et al., 2020).

The iPSCs lines DMD-A and DMD-B were derived from the patient’s skin biopsies, and DMD-C from a blood sample. CTRL-1 was produced from skin biopsies of a healthy male individual and has been characterized previously (Miller et al., 2017; Moreira et al., 2022). Primary myoblasts (pMYO) were kindly provided by Professor Mariz Vainzof from the cell repository of the Human Genome and Stem Cells Research Center (HUG-CELL).

Briefly, fragments from skin biopsies were treated with dispase (1 mg/ml; Sigma-Aldrich) for 24h and migrating fibroblasts were expanded in DMEM High glucose supplemented with 10% fetal bovine serum (FBS) and 100 µg/ml Normocin. Peripheral blood mononuclear cells (PBMC) were obtained through whole blood isolation with FicolPaque (Sigma-Aldrich), expanded in StemSpan medium (StemCell Technologies) supplemented with 50 ng/ml of stem cell factor (SCF), 1 µM dexamethasone, 2U/ml erythropoietin (EPO), 10 ng/ml interleukin 3 (IL-3), 40 ng/ml insulin growth factor-1 (IGF-1), and 100 µg/ml normocin in low attachment plates until ready for reprogramming.

### Reprogramming and characterization of iPSCs

Fibroblasts or PBMCs were reprogrammed using the Human Dermal Fibroblast Nucleofector™ Kit (Lonza) or Human CD34^+^ Cell Nucleofector^TM^ Kit (Lonza), according to the manufacturer’s instructions, with the episomal vectors pCXLE-hOCT3/4-shp53-F, pCXLE-hSK, and pCXLE-hUL (Addgene) using NucleofectorTM 2b Device.

After nucleofection, PBMCs were plated on top of a murine embryonic fibroblast layer in hESC medium (DMEM/F12 supplemented with 2mM GlutaMAX-I, 20% knockout serum replacement (KSR), 1% non-essential amino acids (NEAA), 100 µM 2-mercaptoetanol, 100 µg/ml normocin, 2 µM SB431542, 0.5 µM PD0325901, 2 µM thiazovivin, 0.5 mM valproic acid, 0.25 mM sodium butyrate (NaB), 10 ng/ml fibroblast growth factor 2 (FGF2). For fibroblasts, the same medium was used, but thiazovivin and NaB were omited. The emerging iPSCs colonies were isolated and transferred to Matrigel-coated plates in Essential 8^TM^ medium (Thermo Fisher Scientific) supplemented with 5 µM Y-27632. Colonies were expanded using Essential 8 medium with 100 µg/ml normocin. Characterization and differentiation were performed after at least eight passages.

The expression of pluripotency markers *NANOG*, *SOX2*, and *OCT4* was evaluated using RT-qPCR, and the presence of OCT4 and SSEA4 positive cells was confirmed using immunofluorescence (Supplementary Methods). Multiplex ligation-dependent probe amplification of subtelomeric regions (P036 and P070; MRC-Holland) was performed in iPSC lines as previously described (Jehee et al., 2011) to check for chromosomal imbalances.

### Generation of PAX7 reporter lineages

Using CRISPR-Cas9, we performed a knock-in in exon 1 of *PAX7* promoter in the WT, DMD and DMD-C lineages, as described (Tanoury et al., 2020). Briefly, iPSCs were transfected with plasmids carrying the 2A-NLS-Venus sequence in frame with exon 1 of *PAX7*. Edited cells were selected and genotyped.

### Differentiation into myogenic progenitors: 2D and iSMS culture

Skeletal muscle differentiation was carried out using a transgene-free protocol (Chal et al., 2016) that recapitulates the key molecular events of paraxial mesoderm specification, followed by differentiation into skeletal muscle fibers and progenitors.

Briefly, iPSCs were plated in 6 well Matrigel coated plates (20 µg/cm^2^) at low confluence (2.9 x 10^5^ cells per well) for 2D differentiation. For iSMS, iPSCs were detached using a cell lifter, transferred to 6 well low-attachment plates, and cultivated on an orbital shaker at 65 rpm. This protocol was adapted from a previously described free-floating protocol for deriving EZ spheres from iPSCs (Hosoyama et al., 2014).

In the first two days, the medium was changed to DMEM F12 supplemented with 2 mM GlutaMAX-I, 1% ITS, 1% NEAA, 0.003 mM CHIR-99021, and 0.0005 mM LDN-193189 (Figure 1A, CL medium). On day 3, 20 ng/ml FGF2 was added to the CL medium (CLF). From day 6 to 8, the medium was changed to DMEM High glucose, 2 mM GlutaMAX-I, 15% KSR, 1% NEAA, 0.2% PenStrep, 0.1 mM 2-mercaptoetanol, 0.01 µg/ml of hepatocyte growth factor (HGF), 0.002 µg/ml IGF-1, 0.020 µg/ml FGF2, and 0.0005 mM LDN-193189 (KHIFL medium). The factors HGF, FGF2, and LDN were removed from days 8 to 12 (KI), and the cells were treated with medium supplemented with HGF and IGF from days 12 to 21.

On day 21 of differentiation, cells in the 2D experiment were treated with 0.07% Collagenase IV (ColIV) and TrypLE™ Express Enzyme (1X), filtered through a 70 µM cell strainer and resuspended in myoblast growth medium (SKGM-2, Lonza CC-3245) supplemented with 10 µM Y-27632. The skeletal muscle spheroids (iSMS) generated were maintained in low-attachment plates and the medium was changed to SKGM-2 with 10 µM Y-27632. The cells were maintained in SKGM-2 without Y-27632 for approximately 72 h. To perform secondary differentiation, the medium was changed to DMEM/F12 supplemented with 2 mM GlutaMAX-I, 20% KSR, 1% ITS, 0.2% penicillin streptomycin and 10 µM SB431542.

To check the freezing/thawing capacity, after secondary differentiation, cells from 2D and iSMS were cultivated in SKGM-2 medium. The iSMS were allowed to attach to Matrigel-coated plates and migrate out of the iSMS explants. All myogenic progenitors were frozen in FBS and 10% DMSO.

### Quantitative reverse transcriptase PCR (RT-qPCR)

Total RNA was extracted on day 21 of primary differentiation using QIAzol Lysis Reagent followed by isolation with RNeasy Mini kit (Qiagen), according to the manufacturer’s protocols. WT and DMD were further differentiated (secondary differentiation) and samples were collected on day 8.

About 0.5 µg of RNA was utilized for cDNA synthesis using High-Capacity cDNA Reverse Transcription Kit (ThermoFisher). The cDNA was diluted at a 1/20 ratio and the PCR reaction was set up with primers (3 µM) for *MYOD, MYOG, MYF5, MYH3, PAX7, JAG1, NOTCH1, NOTCH2, HES1, HEYL,* and *NRARP* (Supplementary Table 1) and Fast SYBR^TM^ Green Master Mix. The reactions were run on a QuantStudio™ 12K Flex Real-Time PCR System. Data were processed using Expression Suite Software, fold change was calculated using the ΔΔCt method with WT 2D or CTRL-1 for normalization, and *RPLP0* as endogenous control.

### RT Array – Notch pathway

RNA samples (0.5 µg) from day 21 of differentiation of WT and DMD iPSC lines from 2D and 3D cultures were used for cDNA synthesis with RT^2^ First Strand Kit. Then, master mix with corresponding cDNA was prepared and samples were dispensed in RT^2^ Profiler PCR Array Human Notch Signaling Pathway Plus (PAHS-059YC, Qiagen), according to the manufacturer’s protocol using Applied Biosystems 7500 Fast Real-Time PCR System. Analysis was performed in RT² Profiler PCR Data Analysis platform (https://dataanalysis2.qiagen.com/pcr).

### Immunofluorescence

iSMS on day 21 of differentiation were fixed with 4% paraformaldehyde (PFA) for 1 h. For some stainings, we embedded iSMS in Optimal Cutting Temperature (O.C.T.) compound, immediately frozen in liquid nitrogen and sectioned using a cryostat. For other staining procedures, we performed the protocol using the whole iSMS.

Samples were permeabilized with 0.5% Triton X-100 in phosphate-buffered saline (PBS) for 15 min. Blocking was performed with 30 minutes of incubation with 0.1% Triton X-100 and 10% FBS in PBS. Samples were incubated with primary antibodies against TITIN (DSHB, 1/300), MYOD (ab64159, 1/300), MYF5 (sc302, 1/100), and PAX7 (DSHB, 1/100) overnight at 4°C. Then, incubation for 2-3h was performed with anti-rabbit AlexaFluor 546 or anti-mouse AlexaFluor 488 (1/1000) secondary antibodies and NucBlue^TM^ Fixed Cell Stain ReadyProbes^TM^ reagent (Thermo Fisher Scientific). Immunostained iSMS were analyzed in glass-bottom dishes or mounted slides using a confocal microscope (LSM 800). Confocal Z-stacks were collected and reconstructed into 3D images using ZEN Blue software (Zeiss).

### Flow cytometry

On day 21, PAX7-Venus reporter WT, DMD or DMD-C myogenic cells (from 2D and 3D experiments) were treated with ColIV and TryPLE Express, filtered through 70 µm cell strainer and analyzed on LSRFortessa X-20. Non-reporter cells were used as the negative controls. A total of 50.000 events were acquired.

Non-reporter cells differentiated from iPSCs lines DMD-A, DMD-B, DMD-C, and CTRL-1 were subjected to antibody staining. Briefly, resuspended cells were stained with PAX7 antibody (abcam, 1/50) and anti-rabbit AlexaFluor 546 secondary antibody (1/1000) using Fix & Perm (Life Technologies), according to the manufacturer’s instructions. The stained cells were analyzed using a FACSAria III Cell Sorter. Cells stained with only secondary antibody were used as negative controls. At least 20.000 events were acquired. The median fluorescence intensity (MFI ratio) was calculated by dividing the median fluorescence intensity obtained from stained cells by the median fluorescence intensity from the negative controls. The flow cytometry data were analyzed using FlowJo X software.

### Cell proliferation assay

Myogenic progenitors (single cells) derived from the 2D and iSMS protocols were thawed, expanded in SKGM-2 medium, and plated at a density of 2400 cells/cm^2^. After 4 days, cells were treated with EdU 1 µM for 1 h and detection was performed using the Click-iT® Plus EdU Flow Cytometry Assay Kit, following the manufacturer’s instructions. Cells incubated with EdU only, but not with the reaction cocktail, were used as negative controls. Samples were analyzed using a BD FACSymphony™ A1 Cell Analyzer and at least 20000 events were acquired. FlowJo X software was used for analysis.

### Differentiation of myoblasts from 2D and iSMS cultures

After thawing, the cells were expanded in SKGM-2 medium. About 4000 cells were plated per well in Nunc^®^ Lab-Tek^®^ II Chamber Slides™ (Sigma). When confluence was reached, TDM was added and changed every other day, for eight days. Cells were fixed and immunostained with primary antibodies against TITIN, MYOD, or PAX7 followed by secondary antibodies and nuclei staining, as described in the immunostaining section.

### Statistical Analysis

All experiments were performed in triplicate, except where indicated. For the RT-qPCR analysis of WT and DMD, we employed two-way ANOVA and Bonferroni’s posttest. For the RT-qPCR analysis of patient-derived skeletal muscles, we performed Unpaired t-test, comparing each patient individually (CTRL-1, DMD-A, DMD-B and DMD-C) in 2D versus iSMS. Graphs and statistical analysis were performed using GraphPad Prism version 6.00 for Windows (GraphPad Software, La Jolla, California, USA, www.graphpad.com).

## List of abbreviations

DMD: Duchenne Muscular Dystrophy

SC: Satellite Cells

2D: bidimensional

3D: tridimensional

iSMS: skeletal muscle spheroids

iPSC: induced pluripotent stem cells

RT-qPCR: reverse transcriptase quantitative polymerase chain reaction

YFP: yellow fluorescent protein

WT: wild-type

D21: primary differentiation day 21

SD8: secondary differentiation day 8

pMYO: primary myoblasts

FACS: fluorescence-activated cell sorting

PBMC: peripheral blood mononuclear cells

## Declarations

### Ethics approval and consent to participate

The experiments were approved by the Research Ethics Committee of the Institute of Biosciences at the University of São Paulo (CAAE: 25342719.6.0000.5464) and the patients or their parents signed a consent form.

### Consent for publication

Not applicable.

### Availability of data and materials

All data generated or analyzed during this study are included in this published article.

### Authors’ contributions

JE, FSL, TM and KT: conceptualization; JE, FSL: Writing - Original draft; JE: Data curation, Formal Analysis; JE, FSL, IN, TM, GO, ZAT, KT, MP, RB: methodology, investigation, writing – review and editing; MH: methodology, writing – review and editing. OP, MZ: Methodology and funding acquisition. MZ: Supervision and project administration. All authors have read and approved the final manuscript.

## Acknowledgments

The authors acknowledge the patients and their families; the CEGH-CEL team, especially Patrícia Semedo, Wagner Falciano, Luciana Cristina, Naila Lourenço, Maria Fernanda Amarante, Letícia Nogueira, Marta Rita, Raiane Ferreira, Márcia Pereira, and Mateus Vidigal, and special thanks to Professor Mariz Vainzof for donating cells and resources. We also acknowledge the Research Support Facilities Center (CEFAP-USP).

## REFERENCES

Aisenbrey, E. A., & Murphy, W. L. (2020). Synthetic alternatives to Matrigel. *Nature reviews*. Materials, 5(7), 539–551. 10.1038/s41578-020-0199-8

Ancel, S., Stuelsatz, P., & Feige, J. N. (2021). Muscle Stem Cell Quiescence: Controlling Stemness by Staying Asleep. Trends in Cell Biology, 31(7), 556–568. 10.1016/j.tcb.2021.02.006

Bjornson, C. R. R., Cheung, T. H., Liu, L., Tripathi, P. V., Steeper, K. M., & Rando, T. A. (2012). Notch signaling is necessary to maintain quiescence in adult muscle stem cells. *Stem Cells (Dayton*, Ohio), 30(2), 232–242. 10.1002/stem.773

Blau, H. M., Webster, C., & Pavlath, G. K. (1983). Defective Myoblasts Identified in Duchenne Muscular Dystrophy. Proceedings of the National Academy of Sciences of the United States of America, 80(15), 4856–4860.

Bröhl, D., Vasyutina, E., Czajkowski, M. T., Griger, J., Rassek, C., Rahn, H.-P., Purfürst, B., Wende, H., & Birchmeier, C. (2012). Colonization of the satellite cell niche by skeletal muscle progenitor cells depends on Notch signals. Developmental Cell, 23(3), 469–481. 10.1016/j.devcel.2012.07.014

Chal, J., Al Tanoury, Z., Hestin, M., Gobert, B., Aivio, S., Hick, A., Cherrier, T., Nesmith, A. P., Parker, K. K., & Pourquié, O. (2016). Generation of human muscle fibers and satellite-like cells from human pluripotent stem cells in vitro. Nature Protocols, 11(10), 1833–1850. 10.1038/nprot.2016.110

Chal, J., Oginuma, M., Al Tanoury, Z., Gobert, B., Sumara, O., Hick, A., Bousson, F., Zidouni, Y., Mursch, C., Moncuquet, P., Tassy, O., Vincent, S., Miyanari, A., Bera, A., Garnier, J.-M., Guevara, G., Hestin, M., Kennedy, L., Hayashi, S., … Pourquié, O. (2015). Differentiation of pluripotent stem cells to muscle fiber to model Duchenne muscular dystrophy. Nature Biotechnology, 33(9), 962–969. 10.1038/nbt.3297

Chien, P., Xi, H., & Pyle, A. D. (2022). Recapitulating human myogenesis ex vivo using human pluripotent stem cells. Experimental Cell Research, 411(2), 112990. 10.1016/j.yexcr.2021.112990

Choi, I. Y., Lim, H., Estrellas, K., Mula, J., Cohen, T. V., Zhang, Y., Donnelly, C. J., Richard, J.-P., Kim, Y. J., Kim, H., Kazuki, Y., Oshimura, M., Li, H. L., Hotta, A., Rothstein, J., Maragakis, N., Wagner, K. R., & Lee, G. (2016). Concordant but Varied Phenotypes among Duchenne Muscular Dystrophy Patient-Specific Myoblasts Derived using a Human iPSC-Based Model. Cell Reports, 15(10), 2301–2312. 10.1016/j.celrep.2016.05.016

Danisovic, L., Culenova, M., & Csobonyeiova, M. (2018). Induced Pluripotent Stem Cells for Duchenne Muscular Dystrophy Modeling and Therapy. Cells, 7(12). 10.3390/cells7120253

Dessauge, F., Schleder, C., Perruchot, M.-H., & Rouger, K. (2021). 3D in vitro models of skeletal muscle: Myopshere, myobundle and bioprinted muscle construct. Veterinary Research, 52(1), 72. 10.1186/s13567-021-00942-w

Duan, D., Goemans, N., Takeda, S., Mercuri, E., & Aartsma-Rus, A. (2021). Duchenne muscular dystrophy. Nature Reviews Disease Primers, 7(1), Artigo 1. 10.1038/s41572-021-00248-3

Dumont, N. A., Wang, Y. X., von Maltzahn, J., Pasut, A., Bentzinger, C. F., Brun, C. E., & Rudnicki, M. A. (2015). Dystrophin expression in muscle stem cells regulates their polarity and asymmetric division. Nature Medicine, 21(12), 1455–1463. 10.1038/nm.3990

Emery, A. E. H. (2002). The muscular dystrophies. Lancet (London, England), 359(9307), 687–695. 10.1016/S0140-6736(02)07815-7

Filippelli, R. L., & Chang, N. C. (2021). Empowering Muscle Stem Cells for the Treatment of Duchenne Muscular Dystrophy. Cells Tissues Organs, 1–14. 10.1159/000514305

Gosselin, M. R., Mournetas, V., Borczyk, M., Verma, S., Occhipinti, A., Róg, J., Bozycki, L., Korostynski, M., Robson, S. C., Angione, C., Pinset, C., & Gorecki, D. C. (2022). Loss of full-length dystrophin expression results in major cell-autonomous abnormalities in proliferating myoblasts. eLife, 11, e75521. 10.7554/eLife.75521

Hartog, L. D., & Asakura, A. (2022). Implications of notch signaling in duchenne muscular dystrophy. Frontiers in Physiology, 13, 984373. 10.3389/fphys.2022.984373

Hicks, M. R., Hiserodt, J., Paras, K., Fujiwara, W., Eskin, A., Jan, M., Xi, H., Young, C. S., Evseenko, D., Nelson, S. F., Spencer, M. J., Handel, B. V., & Pyle, A. D. (2018). ERBB3 and NGFR mark a distinct skeletal muscle progenitor cell in human development and hPSCs. Nature Cell Biology, 20(1), 46–57. 10.1038/s41556-017-0010-2

Hoffman, E. P., Brown, R. H., & Kunkel, L. M. (1987). Dystrophin: The protein product of the Duchenne muscular dystrophy locus. Cell, 51(6), 919–928. 10.1016/0092-8674(87)90579-4

Hosoyama, T., McGivern, J. V., Van Dyke, J. M., Ebert, A. D., & Suzuki, M. (2014). Derivation of myogenic progenitors directly from human pluripotent stem cells using a sphere-based culture. Stem Cells Translational Medicine, 3(5), 564–574. 10.5966/sctm.2013-0143

Jalal, S., Dastidar, S., & Tedesco, F. S. (2021). Advanced models of human skeletal muscle differentiation, development and disease: Three-dimensional cultures, organoids and beyond. Current Opinion in Cell Biology, 73, 92–104. 10.1016/j.ceb.2021.06.004

Jehee, F. S., Takamori, J. T., Medeiros, P. F. V., Pordeus, A. C. B., Latini, F. R. M., Bertola, D. R., Kim, C. A., & Passos-Bueno, M. R. (2011). Using a combination of MLPA kits to detect chromosomal imbalances in patients with multiple congenital anomalies and mental retardation is a valuable choice for developing countries. European Journal of Medical Genetics, 54(4), e425–432. 10.1016/j.ejmg.2011.03.007

Jiwlawat, S., Lynch, E., Glaser, J., Smit-Oistad, I., Jeffrey, J., Van Dyke, J. M., & Suzuki, M. (2017). Differentiation and sarcomere formation in skeletal myocytes directly prepared from human induced pluripotent stem cells using a sphere-based culture. Differentiation; Research in Biological Diversity, 96, 70–81. 10.1016/j.diff.2017.07.004

Kawatani, K., Nambara, T., Nawa, N., Yoshimatsu, H., Kusakabe, H., Hirata, K., Tanave, A., Sumiyama, K., Banno, K., Taniguchi, H., Arahori, H., Ozono, K., & Kitabatake, Y. (2021). A human isogenic iPSC-derived cell line panel identifies major regulators of aberrant astrocyte proliferation in Down syndrome. Communications Biology, 4(1), 1–15. 10.1038/s42003-021-02242-7

Khodabukus, A., Prabhu, N., Wang, J., & Bursac, N. (2018). In Vitro Tissue-Engineered Skeletal Muscle Models for Studying Muscle Physiology and Disease. Advanced Healthcare Materials, 7(15), e1701498. 10.1002/adhm.201701498

Kodippili, K., & Rudnicki, M. A. (2023). Satellite cell contribution to disease pathology in Duchenne muscular dystrophy. Frontiers in Physiology, 14, 1180980. 10.3389/fphys.2023.1180980

Kottlors, M., & Kirschner, J. (2010). Elevated satellite cell number in Duchenne muscular dystrophy. Cell and Tissue Research, 340(3), 541–548. 10.1007/s00441-010-0976-6

Lin, S., Shen, H., Jin, B., Gu, Y., Chen, Z., Cao, C., Hu, C., Keller, C., Pear, W. S., & Wu, L. (2013). Brief report: Blockade of Notch signaling in muscle stem cells causes muscular dystrophic phenotype and impaired muscle regeneration. Stem Cells (Dayton, Ohio), 31(4), 823–828. 10.1002/stem.1319

Liu, C., Oikonomopoulos, A., Sayed, N., & Wu, J. C. (2018). Modeling human diseases with induced pluripotent stem cells: From 2D to 3D and beyond. Development (Cambridge, England), 145(5), dev156166. 10.1242/dev.156166

Maffioletti, S. M., Sarcar, S., Henderson, A. B. H., Mannhardt, I., Pinton, L., Moyle, L. A., Steele-Stallard, H., Cappellari, O., Wells, K. E., Ferrari, G., Mitchell, J. S., Tyzack, G. E., Kotiadis, V. N., Khedr, M., Ragazzi, M., Wang, W., Duchen, M. R., Patani, R., Zammit, P. S., … Tedesco, F. S. (2018). Three-Dimensional Human iPSC-Derived Artificial Skeletal Muscles Model Muscular Dystrophies and Enable Multilineage Tissue Engineering. Cell Reports, 23(3), 899–908. 10.1016/j.celrep.2018.03.091

Mavrommatis, L., Jeong, H.-W., Kindler, U., Gomez-Giro, G., Kienitz, M.-C., Stehling, M., Psathaki, O. E., Zeuschner, D., Bixel, M. G., Han, D., Morosan-Puopolo, G., Gerovska, D., Yang, J. H., Kim, J. B., Arauzo-Bravo, M. J., Schwamborn, J. C., Hahn, S. A., Adams, R. H., Schöler, H. R., … Zaehres, H. (2023). Human skeletal muscle organoids model fetal myogenesis and sustain uncommitted PAX7 myogenic progenitors. eLife, 12, RP87081. 10.7554/eLife.87081

Miller, E. E., Kobayashi, G. S., Musso, C. M., Allen, M., Ishiy, F. A. A., de Caires, L. C., Goulart, E., Griesi-Oliveira, K., Zechi-Ceide, R. M., Richieri-Costa, A., Bertola, D. R., Passos-Bueno, M. R., & Silver, D. L. (2017). EIF4A3 deficient human iPSCs and mouse models demonstrate neural crest defects that underlie Richieri-Costa-Pereira syndrome. Human Molecular Genetics, 26(12), 2177–2191. 10.1093/hmg/ddx078

Monaco, A. P., Bertelson, C. J., Liechti-Gallati, S., Moser, H., & Kunkel, L. M. (1988). An explanation for the phenotypic differences between patients bearing partial deletions of the DMD locus. Genomics, 2(1), 90–95. 10.1016/0888-7543(88)90113-9

Montarras, D., Morgan, J., Collins, C., Relaix, F., Zaffran, S., Cumano, A., Partridge, T., & Buckingham, M. (2005). Direct Isolation of Satellite Cells for Skeletal Muscle Regeneration. Science, 309(5743), 2064–2067. 10.1126/science.1114758

Moreira, D. de P., Suzuki, A. M., Silva, A. L. T. e, Varella-Branco, E., Meneghetti, M. C. Z., Kobayashi, G. S., Fogo, M., Ferrari, M. de F. R., Cardoso, R. R., Lourenço, N. C. V., Griesi-Oliveira, K., Zachi, E. C., Bertola, D. R., Weinmann, K. de S., Lima, M. A. de, Nader, H. B., Sertié, A. L., & Passos-Bueno, M. R. (2022). Neuroprogenitor Cells From Patients With TBCK Encephalopathy Suggest Deregulation of Early Secretory Vesicle Transport. Frontiers in Cellular Neuroscience, 15. https://www.frontiersin.org/articles/10.3389/fncel.2021.803302

Mourikis, P., Sambasivan, R., Castel, D., Rocheteau, P., Bizzarro, V., & Tajbakhsh, S. (2012). A critical requirement for notch signaling in maintenance of the quiescent skeletal muscle stem cell state. Stem Cells (Dayton, Ohio), 30(2), 243–252. 10.1002/stem.775

Nguyen, H. N., Byers, B., Cord, B., Shcheglovitov, A., Byrne, J., Gujar, P., Kee, K., Schüle, B., Dolmetsch, R. E., Langston, W., Palmer, T. D., & Pera, R. R. (2011). LRRK2 mutant iPSC-derived DA neurons demonstrate increased susceptibility to oxidative stress. Cell Stem Cell, 8(3), 267–280. 10.1016/j.stem.2011.01.013

Oliveira, D., Morales-Vicente, D. A., Amaral, M. S., Luz, L., Sertié, A. L., Leite, F. S., Navarro, C., Kaid, C., Esposito, J., Goulart, E., Caires, L., Alves, L. M., Melo, U. S., Figueiredo, T., Mitne-Neto, M., Okamoto, O. K., Verjovski-Almeida, S., & Zatz, M. (2020). Different gene expression profiles in iPSC-derived motor neurons from ALS8 patients with variable clinical courses suggest mitigating pathways for neurodegeneration. Human Molecular Genetics, 29(9), 1465–1475. 10.1093/hmg/ddaa069

Pang, K. T., Loo, L. S. W., Chia, S., Ong, F. Y. T., Yu, H., & Walsh, I. (2023). Insight into muscle stem cell regeneration and mechanobiology. Stem Cell Research & Therapy, 14(1), 129. 10.1186/s13287-023-03363-y

Picard, C. A., & Marcelle, C. (2013). Two distinct muscle progenitor populations coexist throughout amniote development. Developmental Biology, 373(1), 141–148. 10.1016/j.ydbio.2012.10.018

Piga, D., Salani, S., Magri, F., Brusa, R., Mauri, E., Comi, G. P., Bresolin, N., & Corti, S. (2019). Human induced pluripotent stem cell models for the study and treatment of Duchenne and Becker muscular dystrophies. Therapeutic Advances in Neurological Disorders, 12. 10.1177/1756286419833478

Rao, L., Qian, Y., Khodabukus, A., Ribar, T., & Bursac, N. (2018). Engineering human pluripotent stem cells into a functional skeletal muscle tissue. Nature Communications, 9(1), Artigo 1. 10.1038/s41467-017-02636-4

Ribeiro, A. F., Souza, L. S., Almeida, C. F., Ishiba, R., Fernandes, S. A., Guerrieri, D. A., Santos, A. L. F., Onofre-Oliveira, P. C. G., & Vainzof, M. (2019). Muscle satellite cells and impaired late stage regeneration in different murine models for muscular dystrophies. Scientific Reports, 9(1), Artigo 1. 10.1038/s41598-019-48156-7

Sacco, A., Mourkioti, F., Tran, R., Choi, J., Llewellyn, M., Kraft, P., Shkreli, M., Delp, S., Pomerantz, J. H., Artandi, S. E., & Blau, H. M. (2010). Short Telomeres and Stem Cell Exhaustion Model Duchenne Muscular Dystrophy in mdx/mTR Mice. Cell, 143(7), 1059–1071. 10.1016/j.cell.2010.11.039

Sakai-Takemura, F., Narita, A., Masuda, S., Wakamatsu, T., Watanabe, N., Nishiyama, T., Nogami, K., Blanc, M., Takeda, S., & Miyagoe-Suzuki, Y. (2018). Premyogenic progenitors derived from human pluripotent stem cells expand in floating culture and differentiate into transplantable myogenic progenitors. Scientific Reports, 8(1), Artigo 1. 10.1038/s41598-018-24959-y

Seale, P., Sabourin, L. A., Girgis-Gabardo, A., Mansouri, A., Gruss, P., & Rudnicki, M. A. (2000). Pax7 is required for the specification of myogenic satellite cells. Cell, 102(6), 777–786. 10.1016/s0092-8674(00)00066-0

Servián-Morilla, E., Takeuchi, H., Lee, T. V., Clarimon, J., Mavillard, F., Area-Gómez, E., Rivas, E., Nieto-González, J. L., Rivero, M. C., Cabrera-Serrano, M., Gómez-Sánchez, L., Martínez-López, J. A., Estrada, B., Márquez, C., Morgado, Y., Suárez-Calvet, X., Pita, G., Bigot, A., Gallardo, E., … Paradas, C. (2016). A POGLUT1 mutation causes a muscular dystrophy with reduced Notch signaling and satellite cell loss. EMBO Molecular Medicine, 8(11), 1289–1309. 10.15252/emmm.201505815

Shin, M.-K., Bang, J. S., Lee, J. E., Tran, H.-D., Park, G., Lee, D. R., & Jo, J. (2022). Generation of Skeletal Muscle Organoids from Human Pluripotent Stem Cells to Model Myogenesis and Muscle Regeneration. International Journal of Molecular Sciences, 23(9), 5108. 10.3390/ijms23095108

Shoji, E., Sakurai, H., Nishino, T., Nakahata, T., Heike, T., Awaya, T., Fujii, N., Manabe, Y., Matsuo, M., & Sehara-Fujisawa, A. (2015). Early pathogenesis of Duchenne muscular dystrophy modelled in patient-derived human induced pluripotent stem cells. Scientific Reports, 5(1), Artigo 1. 10.1038/srep12831

Spinazzola, J. M., & Gussoni, E. (2017). Isolation of Primary Human Skeletal Muscle Cells. Bio-protocol, 7(21), e2591. 10.21769/BioProtoc.2591

Takahashi, K., Tanabe, K., Ohnuki, M., Narita, M., Ichisaka, T., Tomoda, K., & Yamanaka, S. (2007). Induction of pluripotent stem cells from adult human fibroblasts by defined factors. Cell, 131(5), 861–872. 10.1016/j.cell.2007.11.019

Tanoury, Z. A., Rao, J., Tassy, O., Gobert, B., Gapon, S., Garnier, J.-M., Wagner, E., Hick, A., Hall, A., Gussoni, E., & Pourquié, O. (2020). Differentiation of the human PAX7-positive myogenic precursors/satellite cell lineage in vitro. Development, 147(12). 10.1242/dev.187344

Tanoury, Z. A., Zimmerman, J. F., Rao, J., Sieiro, D., McNamara, H. M., Cherrier, T., Rodríguez-delaRosa, A., Hick-Colin, A., Bousson, F., Fugier-Schmucker, C., Marchiano, F., Habermann, B., Chal, J., Nesmith, A. P., Gapon, S., Wagner, E., Gupta, V. A., Bassel-Duby, R., Olson, E. N., … Pourquié, O. (2021). Prednisolone rescues Duchenne muscular dystrophy phenotypes in human pluripotent stem cell–derived skeletal muscle in vitro. Proceedings of the National Academy of Sciences, 118(28). 10.1073/pnas.2022960118

Tierney, M. T., & Sacco, A. (2016). Satellite Cell Heterogeneity in Skeletal Muscle Homeostasis. Trends in Cell Biology, 26(6), 434–444. 10.1016/j.tcb.2016.02.004

Vieira, N. M., Elvers, I., Alexander, M. S., Moreira, Y. B., Eran, A., Gomes, J. P., Marshall, J. L., Karlsson, E. K., Verjovski-Almeida, S., Lindblad-Toh, K., Kunkel, L. M., & Zatz, M. (2015). Jagged 1 Rescues the Duchenne Muscular Dystrophy Phenotype. Cell, 163(5), 1204–1213. 10.1016/j.cell.2015.10.049

Webster, C., & Blau, H. M. (1990). Accelerated age-related decline in replicative life-span of Duchenne muscular dystrophy myoblasts: Implications for cell and gene therapy. Somatic Cell and Molecular Genetics, 16(6), 557–565. 10.1007/BF01233096

Wells, D. J. (2018). Tracking progress: An update on animal models for Duchenne muscular dystrophy. Disease Models & Mechanisms, 11(6), dmm035774. 10.1242/dmm.035774

Wen, Y., Bi, P., Liu, W., Asakura, A., Keller, C., & Kuang, S. (2012). Constitutive Notch activation upregulates Pax7 and promotes the self-renewal of skeletal muscle satellite cells. Molecular and Cellular Biology, 32(12), 2300–2311. 10.1128/MCB.06753-11

Westerman, K. A., Penvose, A., Yang, Z., Allen, P. D., & Vacanti, C. A. (2010). Adult muscle “stem” cells can be sustained in culture as free-floating myospheres. Experimental Cell Research, 316(12), 1966–1976. 10.1016/j.yexcr.2010.03.022

Xi, H., Langerman, J., Sabri, S., Chien, P., Young, C. S., Younesi, S., Hicks, M., Gonzalez, K., Fujiwara, W., Marzi, J., Liebscher, S., Spencer, M., Van Handel, B., Evseenko, D., Schenke-Layland, K., Plath, K., & Pyle, A. D. (2020). A Human Skeletal Muscle Atlas Identifies the Trajectories of Stem and Progenitor Cells across Development and from Human Pluripotent Stem Cells. Cell Stem Cell, 27(1), 158–176.e10. 10.1016/j.stem.2020.04.017

Yagi, T., Ito, D., Okada, Y., Akamatsu, W., Nihei, Y., Yoshizaki, T., Yamanaka, S., Okano, H., & Suzuki, N. (2011). Modeling familial Alzheimer’s disease with induced pluripotent stem cells. Human Molecular Genetics, 20(23), 4530–4539. 10.1093/hmg/ddr394

Zammit, P. S. (2017). Function of the myogenic regulatory factors Myf5, MyoD, Myogenin and MRF4 in skeletal muscle, satellite cells and regenerative myogenesis. Seminars in Cell & Developmental Biology, 72, 19–32. 10.1016/j.semcdb.2017.11.011

Zhou B., B., Lin, W., Long, Y., Yang, Y., Zhang, H., Wu, K., & Chu, Q. (2022). Notch signaling pathway: Architecture, disease, and therapeutics. Signal Transduction and Targeted Therapy, 7(1), Artigo 1. 10.1038/s41392-022-00934-y

Zhou, S., Han, L., & Wu, Z. (2022). A Long Journey before Cycling: Regulation of Quiescence Exit in Adult Muscle Satellite Cells. International Journal of Molecular Sciences, 23(3), Artigo 3. 10.3390/ijms23031748

Zschüntzsch, J., Meyer, S., Shahriyari, M., Kummer, K., Schmidt, M., Kummer, S., & Tiburcy, M. (2022). The Evolution of Complex Muscle Cell In Vitro Models to Study Pathomechanisms and Drug Development of Neuromuscular Disease. Cells, 11(7), 1233. 10.3390/cells11071233

