## Supplemental Material for "iPSC-derived skeletal muscle spheroids for Duchenne Muscular Dystrophy modeling"

### Supplementary Material

#### Supplementary Methods

1. *Immunostaining of iPSCs*

iPSCs were fixed with 10% formalin for 20 min, permeabilized with 0.2% Triton X-100 in PBS for 30 min and blocked with 5% BSA solution for 1 h. They were then incubated with primary antibodies against SSEA4 and OCT3/4 *overnight* at 4°C, washed with PBS, and treated with AlexaFluor anti-mouse 546 e AlexaFluor anti-rabbit 488 secondary antibodies (1/1000) for 1 h and 4′,6-diamidino-2-phenylindole (DAPI) for 2 min. Images were acquired using confocal microscope LSM 800 (Zeiss) with ZEN software.

2. *Primer sequences*

Supplementary Table 1. Primer sequences for RT-qPCR experiments.

| Refseq | Gene name | Sequence |
|---|---|---|
| **NM_001357943.2** | *GAPDH* | F: TGGTATCGTGGAAGGACTCATG |
| | | R: AGAGGCAGGGATGATGTTCTG |
| **NM_024865.4** | *NANOG* | F: CAGAAGGCCTCAGCACCTAC |
| | | R: ATTGTTCCAGGTCTGGTTGC |
| **NM_003106.4** | *SOX2* | F: TGGGTTCGGTGGTCAAGTCC |
| | | R: CTGGAGTGGGAGGAAGAGGTAAC |
| **NM_001285986.2** | *OCT4* | F: ATGTGGTCCGAGTGTGGTTC |
| | | R: GACCCAGCAGCCTCAAAATCC |
| **NM_002478.5** | *MYOD1* | F: GCCACAACGGACGACTTCTATG |
| | | R: TGCTCTTCGGGTTTCAGGAG |
| **NM_002470.4** | *MYH3* | F: ACCTGAAGGACCGTTACACATC |
| | | R: GCCACTTGTAGGGGTTGACA |
| **NM_013945.3** | *PAX7* | F: AGAAGGCCAAACACAGCATC |
| | | R: TCAGGTTCCGACTCCACATC |
| **NM_005593.3** | *MYF5* | F: ATGCCATCCGCTACATCGAG |
| | | R: CAGACAGGACTGTTACATTCGG |
| **NM_002479.6** | *MYOG* | F: AGTGCCATCCAGTACATCGAG |
| | | R: TGTGAGAGCTGCATTCGC |
| **NM_002479.6** | *RPLP0* | F: AATCTCCAGGGGCACCATTG |
| | | R: GAACACCTGCTGGATGACC |
| **NM_000214.3** | *JAG1* | F: GAAGTAAGAGTTCAGAGGCGGC |
| | | R: GTCACCAAGCAACAGATCCAAG |

| | | |
|---|---|---|
| **NM_017617.5** | *NOTCH1* | F: CGAGTCCGTCATCAATGGCT |
| | | R: GCAGGTACGAGCGTCATTCT |
| **NM_024408.4** | *NOTCH2* | F: CCAATGCCCAGGACAACATG |
| | | R: CAGTTCTGCCACCATTCCC |
| **NM_005524.4** | *HES1* | F: GCTGGAGAAGGCGGACATTC |
| | | R: GGTCACCTCGTTCATGCACT |
| **NM_014571.4** | *HEYL* | F: TGACGGTGGATCACTTGAAA |
| | | R: AGCTGTTGAGGTGGGAGAGA |
| **NM_001004354.3** | *NRARP* | F: AGAACATGACCAACTGCGAGT |
| | | R: TTGACCAGCAGCTTCACGAG |

*3. Primary myoblasts (pMYO) culture, cell sorting and differentiation*

The pMYO from a healthy individual were thawed and expanded in Matrigel-coated plates in SKGM-2 medium. Cells were treated with TryPLE Express, resuspended in 2% FBS in PBS and stained with CD56 and CD82 antibodies, as previously described (Spinazzola & Gussoni, 2017). Cell sorting was performed using BD FACSAria II Cell Sorter. The $CD56^+/CD82^+$ cells were expanded and differentiated for 5 days using TDM. RNA was collected as described in the Methods section.

### Supplementary Figures

#### Supplementary Figure 1

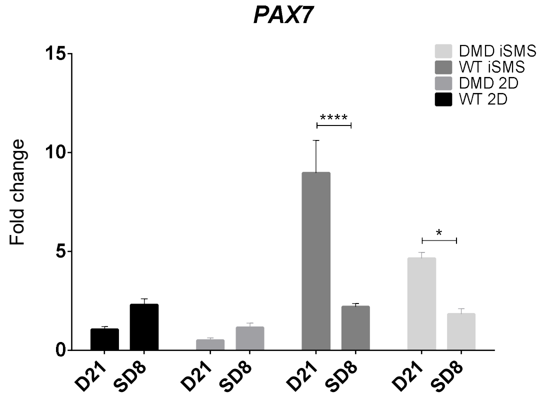


Additional analysis of *PAX7* mRNA expression comparing day 21 of primary differentiation (D21) with day 8 of secondary differentiation (SD8) in WT and DMD. WT 2D at D21 was used for normalization using the ΔΔCT method. TwoWay ANOVA and Sidak's post-test comparing D21 with SD8 was performed. * represents $p<0.05$, and **** represents $p<0.0001$.

#### Supplementary Figure 2

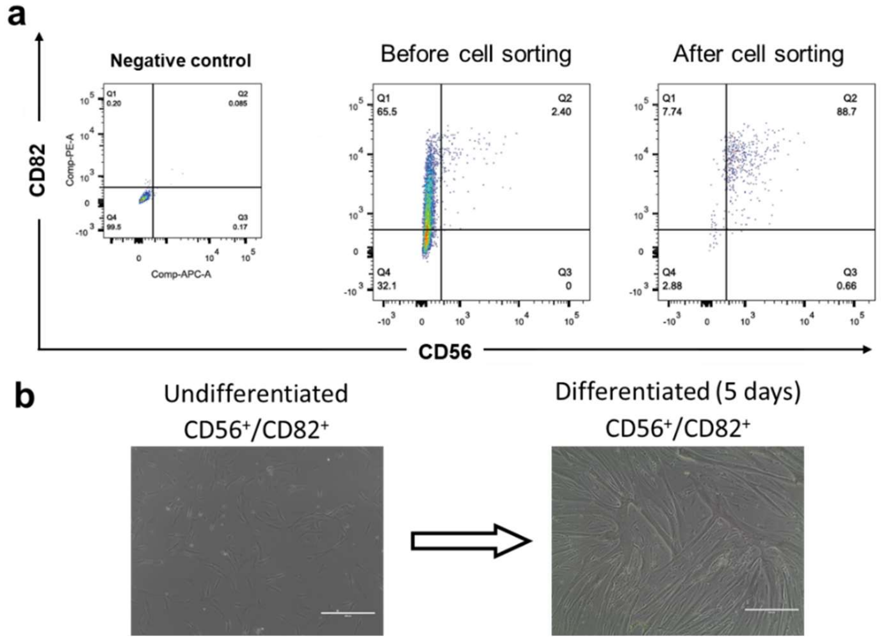


Human primary myoblasts (pMYO) purification through cell sorting.

a. pMYO were FACS sorted using CD56 and CD82 membrane markers to enrich the myogenic population. b. Representative images of undifferentiated $CD56^{+}/CD82^{+}$ pMYO cell and 5 days after myogenic differentiation using TDM medium.

**Supplementary Figure 3**

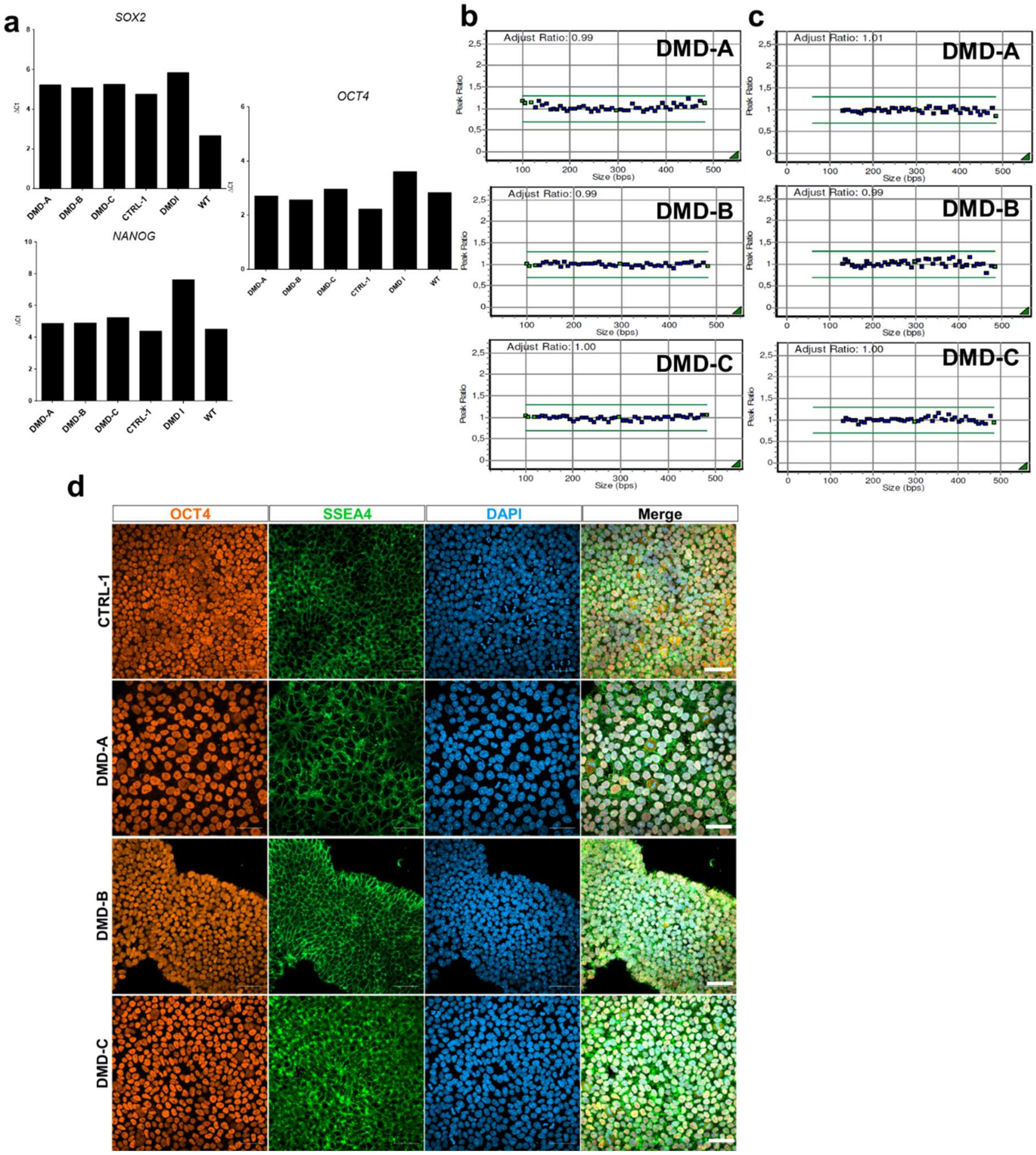


Characterization of iPSC from DMD patients.

a. qPCR of pluripotent stem cell markers *SOX2*, *OCT4*, and *NANOG*. Gene expression data were calculated by ΔCT using *GAPDH* as endogenous control. b and c. Multiplex ligation-dependent probe amplification analysis showing the verification of genomic imbalances in the iPSCs produced. The probes used were P036 (b) and P070 (c) (MRC-Holland). d. Immunofluorescence showing OCT4 (orange) and SSEA4 expression. Scale: 50 µm.

Supplementary Table 2. Description of the iPSC lines from DMD patients produced in this study and the corresponding *DMD* mutations.

| iPSC line | Cell source | *DMD* mutation |
|---|---|---|
| DMD-A | Skin fibroblasts | Exon 51 to 54 deletion |
| DMD-B | Skin fibroblasts | Point mutation at intron 22 (c.2949+1G>A) |
| DMD-C | PBMC | Exon 17 duplication (out of frame) |

Supplementary Table 3. Genes of the $RT^2$ profiler array for Notch pathway. Genes that were not detected in our samples are shown in red.

| Refseq | Symbol | Description |
|---|---|---|
| NM_001110 | ADAM10 | ADAM metallopeptidase domain 10 |
| NM_003183 | ADAM17 | ADAM metallopeptidase domain 17 |
| NM_001130 | AES | Amino-terminal enhancer of split |
| NM_003502 | AXIN1 | Axin 1 |
| NM_005188 | CBL | Cas-Br-M (murine) ecotropic retroviral transforming sequence |
| NM_053056 | CCND1 | Cyclin D1 |
| NM_001238 | CCNE1 | Cyclin E1 |
| NM_000610 | CD44 | CD44 molecule (Indian blood group) |
| NM_000389 | CDKN1A | Cyclin-dependent kinase inhibitor 1A (p21, Cip1) |
| NM_003879 | CFLAR | CASP8 and FADD-like apoptosis regulator |
| NM_001278 | CHUK | Conserved helix-loop-helix ubiquitous kinase |
| NM_001904 | CTNNB1 | Catenin (cadherin-associated protein), beta 1, 88kDa |
| NM_005618 | DLL1 | Delta-like 1 (Drosophila) |
| NM_016941 | DLL3 | Delta-like 3 (Drosophila) |
| NM_019074 | DLL4 | Delta-like 4 (Drosophila) |
| NM_001429 | EP300 | E1A binding protein p300 |
| NM_004448 | ERBB2 | V-erb-b2 erythroblastic leukemia viral oncogene homolog 2, neuro/glioblastoma derived oncogene homolog (avian) |
| NM_004469 | VEGFD | Vascular endothelial growth factor D |
| NM_005252 | FOS | FBJ murine osteosarcoma viral oncogene homolog |
| NM_005438 | FOSL1 | FOS-like antigen 1 |
| NM_003507 | FZD7 | Frizzled family receptor 7 |
| NM_005269 | GLI1 | GLI family zinc finger 1 |
| NM_002093 | GSK3B | Glycogen synthase kinase 3 beta |
| NM_004964 | HDAC1 | Histone deacetylase 1 |
| NM_005524 | HES1 | Hairy and enhancer of split 1, (Drosophila) |
| NM_012258 | HEY1 | Hairy/enhancer-of-split related with YRPW motif 1 |
| NM_024015 | HOXB4 | Homeobox B4 |

| | | |
|---|---|---|
| NM_018411 | HR | Hairless homolog (mouse) |
| NM_002165 | ID1 | Inhibitor of DNA binding 1, dominant negative helix-loop-helix protein |
| NM_000619 | IFNG | Interferon, gamma |
| NM_000417 | IL2RA | Interleukin 2 receptor, alpha |
| NM_000214 | JAG1 | Jagged 1 |
| NM_002226 | JAG2 | Jagged 2 |
| NM_006121 | KRT1 | Keratin 1 |
| NM_001040167 | LFNG | LFNG O-fucosylpeptide 3-beta-N-acetylglucosaminyltransferase |
| NM_005574 | LMO2 | LIM domain only 2 (rhombotin-like 1) |
| NM_000427 | LOR | Loricrin |
| NM_002335 | LRP5 | Low density lipoprotein receptor-related protein 5 |
| NM_014757 | MAML1 | Mastermind-like 1 (Drosophila) |
| NM_032427 | MAML2 | Mastermind-like 2 (Drosophila) |
| NM_002405 | MFNG | MFNG O-fucosylpeptide 3-beta-N-acetylglucosaminyltransferase |
| NM_002423 | MMP7 | Matrix metallopeptidase 7 (matrilysin, uterine) |
| NM_006312 | NCOR2 | Nuclear receptor corepressor 2 |
| NM_015331 | NCSTN | Nicastrin |
| NM_003998 | NFKB1 | Nuclear factor of kappa light polypeptide gene enhancer in B-cells 1 |
| NM_002502 | NFKB2 | Nuclear factor of kappa light polypeptide gene enhancer in B-cells 2 (p49/p100) |
| NM_017617 | NOTCH1 | Notch 1 |
| NM_024408 | NOTCH2 | Notch 2 |
| NM_000435 | NOTCH3 | Notch 3 |
| NM_004557 | NOTCH4 | Notch 4 |
| NM_003744 | NUMB | Numb homolog (Drosophila) |
| NM_016734 | PAX5 | Paired box 5 |
| NM_172236 | POFUT1 | Protein O-fucosyltransferase 1 |
| NM_015869 | PPARG | Peroxisome proliferator-activated receptor gamma |
| NM_000021 | PSEN1 | Presenilin 1 |
| NM_000447 | PSEN2 | Presenilin 2 (Alzheimer disease 4) |
| NM_172341 | PSENEN | Presenilin enhancer 2 homolog (C. elegans) |
| NM_138296 | PTCRA | Pre T-cell antigen receptor alpha |
| NM_014276 | RBPJL | Recombination signal binding protein for immunoglobulin kappa J region-like |
| NM_002917 | RFNG | RFNG O-fucosylpeptide 3-beta-N-acetylglucosaminyltransferase |
| NM_001754 | RUNX1 | Runt-related transcription factor 1 |
| NM_005065 | SEL1L | Sel-1 suppressor of lin-12-like (C. elegans) |
| NM_000193 | SHH | Sonic hedgehog |
| NM_005631 | SMO | Smoothened, frizzled family receptor |
| NM_012245 | SNW1 | SNW domain containing 1 |
| NM_003153 | STAT6 | Signal transducer and activator of transcription 6, interleukin-4 induced |
| NM_005077 | TLE1 | Transducin-like enhancer of split 1 (E(sp1) homolog, Drosophila) |

| | | |
|---|---|---|
| NM_001204869 | CCN4 | Cellular communication network factor 4 |
| NM_000022 | ADA | Adenosine deaminase |
| NM_032550 | AFAP1L2 | Actin filament associated protein 1-like 2 |
| NM_004416 | DTX1 | Deltex homolog 1 (Drosophila) |
| NR_002196 | H19 | H19, imprinted maternally expressed transcript (non-protein coding) |
| NM_021170 | Hes4 | Hairy and enhancer of split 4 (Drosophila) |
| NM_001010926 | HES5 | Hairy and enhancer of split 5 (Drosophila) |
| NM_012259 | HEY2 | Hairy/enhancer-of-split related with YRPW motif 2 |
| NM_014571 | HEYL | Hairy/enhancer-of-split related with YRPW motif-like |
| NM_001004354 | NRARP | Notch-regulated ankyrin repeat protein |
| NM_003357 | SCGB1A1 | Secretoglobin, family 1A, member 1 (uteroglobin) |
| NM_001085 | SERPINA3 | Serpin peptidase inhibitor, clade A (alpha-1 antiproteinase, antitrypsin), member 3 |
| NM_003044 | SLC6A12 | Solute carrier family 6 (neurotransmitter transporter, betaine/GABA), member 12 |
| NM_003068 | SNAI2 | Snail homolog 2 (Drosophila) |
| NM_003225 | TFF1 | Trefoil factor 1 |
| NM_003810 | TNFSF10 | Tumor necrosis factor (ligand) superfamily, member 10 |
| NM_006398 | UBD | Ubiquitin D |
| NM_001101 | ACTB | Actin, beta |
| NM_004048 | B2M | Beta-2-microglobulin |
| NM_002046 | GAPDH | Glyceraldehyde-3-phosphate dehydrogenase |
| NM_000194 | HPRT1 | Hypoxanthine phosphoribosyltransferase 1 |
| NM_001002 | RPLP0 | Ribosomal protein, large, P0 |
